# The double-edged role of FASII regulator FabT in *Streptococcus pyogenes* infection

**DOI:** 10.1101/2023.06.07.543851

**Authors:** Clara Lambert, Marine Gaillard, Paprapach Wongdontree, Caroline Bachmann, Antoine Hautcoeur, Karine Gloux, Thomas Guilbert, Celine Méhats, Bastien Prost, Audrey Solgadi, Sonia Abreu, Muriel Andrieu, Claire Poyart, Alexandra Gruss, Agnes Fouet

## Abstract

In *Streptococcus pyogenes*, the type II fatty acid (FA) synthesis pathway FASII is feedback-controlled by the FabT repressor bound to an acyl-Acyl carrier protein. Despite FabT defects being linked to reduced virulence in animal models, spontaneous *fabT* mutants arise *in vivo*. To resolve this paradox, we characterized the conditions and mechanisms that require FabT activity, and those that promote *fabT* mutant emergence. The primary *fabT* mutant defect is energy dissipation: specific nutrients are consumed, but the mutant fails to grow on human tissue, cells, or cell filtrates where nutrients are limited. These features explain the FabT requirement during infection. Conversely, *fabT* mutants exhibited a marked growth advantage over the wild-type in biotopes rich in saturated FAs. *fabT* mutants emerge in this context, where continued FASII activity prevented environmental FA incorporation. An *ex vivo* muscle model demonstrated that wild-type *S. pyogenes* was inhibited, while *fabT* mutant growth was stimulated, but conditional to FASII activity. Our findings elucidate the rationale for emerging *fabT* mutants that improve survival in lipid-rich biotopes, but lead to a genetic impasse for infection.

## Introduction

Bacterial membranes form a mutable permeable barrier that facilitates adaptation to a changing environment. They usually comprise a phospholipid bilayer composed of a polar head and apolar fatty acid (FA) chains. The FA synthesis pathway (FASII), which is widespread among bacteria, synthesizes saturated and/or unsaturated FAs joined to an acyl carrier protein (ACP; forming acyl-ACP) (Supplementary Fig. 1a). Unsaturated FAs are produced *via* a FASII shunt catalyzed by an acyl-ACP-isomerase named FabM in streptococci. While FASII is conserved (with some enzyme variation), its regulators differ among Firmicutes. In streptococcaceae and enterococcaccae, FASII gene expression is controlled by a unique MarR-family feedback-type regulator named FabT encoded in the main FASII locus (Supplementary Fig. 1 b-c). FabT uses acyl-ACP as corepressor [^1,2^, for review ^3^]. The affinity of FabT-(acyl-ACP) binding to a specific DNA palindromic sequence increases with the length of the acyl carbon chain and the presence of an unsaturation ^4^. FabT regulons were characterized in various streptococcaceae species and in conditions that affect membrane FA composition, including growth temperature, pH or growth phase ^5–9^. FabT exerts greater repression of genes encoding elongation steps, mainly the *trans*-2-enoyl-ACP reductase II FabK, and less repression of *fabM*. In *Streptococcus pneumoniae fabM* expression is not repressed by FabT ^6^. Accordingly, an *S. pneumoniae* strain lacking a functional FabT produced longer and more saturated FAs ^6^. FabT regulons reportedly also comprise non-FASII genes involved in transport, DNA and carbohydrate metabolism, protein, purine and pyrimidine synthesis; however, their identities vary according to reports and the species under study ^3^. To date, FabT regulons were not analyzed in the presence of exogeneous FAs (eFAs), which enhance FabT transcriptional repression ^4^. This missing information is particularly relevant as numerous host infection sites are FA-rich.

*Streptococcus pyogenes*, also known as Group A *Streptococcus*, GAS, is a major human pathogen causing mainly extracellular infections, which is responsible for a large variety of clinical manifestations ranging from superficial infections to life-threatening invasive infections ^10^. GAS infections rank among the top ten causes of death due to bacterial infections worldwide ^11^. GAS isolates mutated in *fabT* were recovered in non-human primates at the point of intramuscular inoculation, raising the possibility of such populations forming in the human host ^5^. In a murine model, strains harboring *fabT* point mutations display smaller size lesions, no loss of body weight and a lower mortality than their wild-type counterparts ^12^. In a non-human primate model, a *fabT* deleted strain shows decreased colonization and dissemination capacities compared to the parental strain ^5^. Its survival is also decreased in human blood or in the presence of human polymorphonuclear leukocytes ^5^.

These reported properties indicate that the *fabT* mutant variants are poorly adapted for infection, which led us to question the rationale for their emergence. We solve this question here by performing in-depth analyses of the features of WT and a representative *fabT* mutant in different conditions relevant to host infection. We report that the *fabT* mutant is metabolically wasteful, by consuming sugars and amino acids, yet failing to grow, which accounts for its failure to cause infection. Conversely, saturated FA-rich environments impose a counter-selective pressure against WT bacteria expressing active FabT. We show that *fabT* mutant growth is stimulated around lipid-rich muscle sources in a FASII-dependent manner, while WT growth is inhibited. These findings solve the apparent contradiction between the *in vivo* emergence of attenuated *fabT* variants and the need for an active FabT repressor during infection.

## Results

### WT and *fabT* mutant growth properties

Multiple independent FabT point mutants were harvested from the infection site of nonhuman primates, some of which mapped to His105 ^5^. We chose the FabT^H105Y^ point mutation as being representative of mutations that arose *in vivo*, which was established in the *emm28* reference strain M28PF1 (respectively mFabT and WT) ^13,14^ (Supplementary Fig. 1b-c) ^15^. A Δ*fabT* deletion strain was constructed for phenotypic comparisons (Supplementary Fig. 2). mFabT and WT strains grow similarly in laboratory medium (THY), and in THY supplemented with 0.1 % Tween 80 (THY-Tween, a C18:1Δ9 [oleic acid] source) (Supplementary Fig. 2a-b, left; also see ^2^). Survival of WT and mFabT strains in mid-exponential phase was also comparable, as assessed by live-dead staining (Supplementary Fig. 2a-b, right). Unlike mFabT, the Δ*fabT* mutant grew slowly in both THY and THY-Tween media (Supplementary Fig. 2c-d; also see ^12^). As *fabT* deletions were not reported to arise *in vivo,* we chose to study the mFabT mutant strain as being representative of *in vivo* mutations.

## FabT^H105Y^ impacts membrane lipid production and species

The GAS mFabT strain produced greater proportions of longer length saturated FA (C18:0) than the WT strain (26.1 % *versus* 6.8 % respectively; Supplementary Table 1), as first reported in *S. pneumoniae* ^6^. In THY-Tween, which supplies C18:1Δ9, the native FabT represses FASII, and contains 1.7 times more C18:1Δ9 than does mFabT. In this condition, the proportion of C18:0 remained higher in mFabT than in the WT (12.4 % and 0.2 % respectively; Supplementary Table 1) ^2^. For comparison, the Δ*fabT* mutant showed a higher C18:0 to C16:0 ratio than mFabT in both THY and THY-Tween, likely reflecting complete FASII derepression in the deletion strain (Supplementary Table 1). The more extreme membrane FA differences may account for poor Δ*fabT* growth in laboratory media.

The effects of the FabT mutation on membrane FA composition should also alter phospholipid metabolism. Lipid analyses of WT and mFabT strains cultured in THY and THY- Tween media identified notable differences in membrane lipid features: i, the primary differences between WT and mFabT FA composition were reflected in the identifiable lipid species (Supplementary Fig. 3, Supplementary Table 2); ii, THY grown mFabT has ∼60 % overall lipid yield compared to that of the WT, extracted from equivalent bacterial OD600 (N=4); these differences were narrowed to ∼5 % in cultures grown in THY-Tween (Supplementary Table 3). We cannot explain these differences in yield, but suggest that the different FA composition of mFabT might affect extraction efficiency. The reasons for these differences in yield will require further study. Among the detected lipids, diglucosyldiacylglycerol (DGDG) and cardiolipin (CL) amounts were proportionately lower in mFabT compared to WT in THY medium. In THY Tween, levels of both CL and DGDG decreased for WT, but not for mFabT (Fig. 1a-b). iii, CL species in mFabT were 2-fold enriched in deoxidized cardiolipin (deoxy- CL) species compared to CL in the WT strain, regardless of the growth medium (Fig. 1c-e). CL but not deoxy-CL reportedly facilitates oxidative phosphorylation activity by binding protons^16^. In conclusion, the greater proportions of saturated and longer FAs in the *fabT* mutant appear to cause a reduced overall membrane lipid content and alterations in lipid distribution and composition as compared to the WT.

**Fig. 1.**
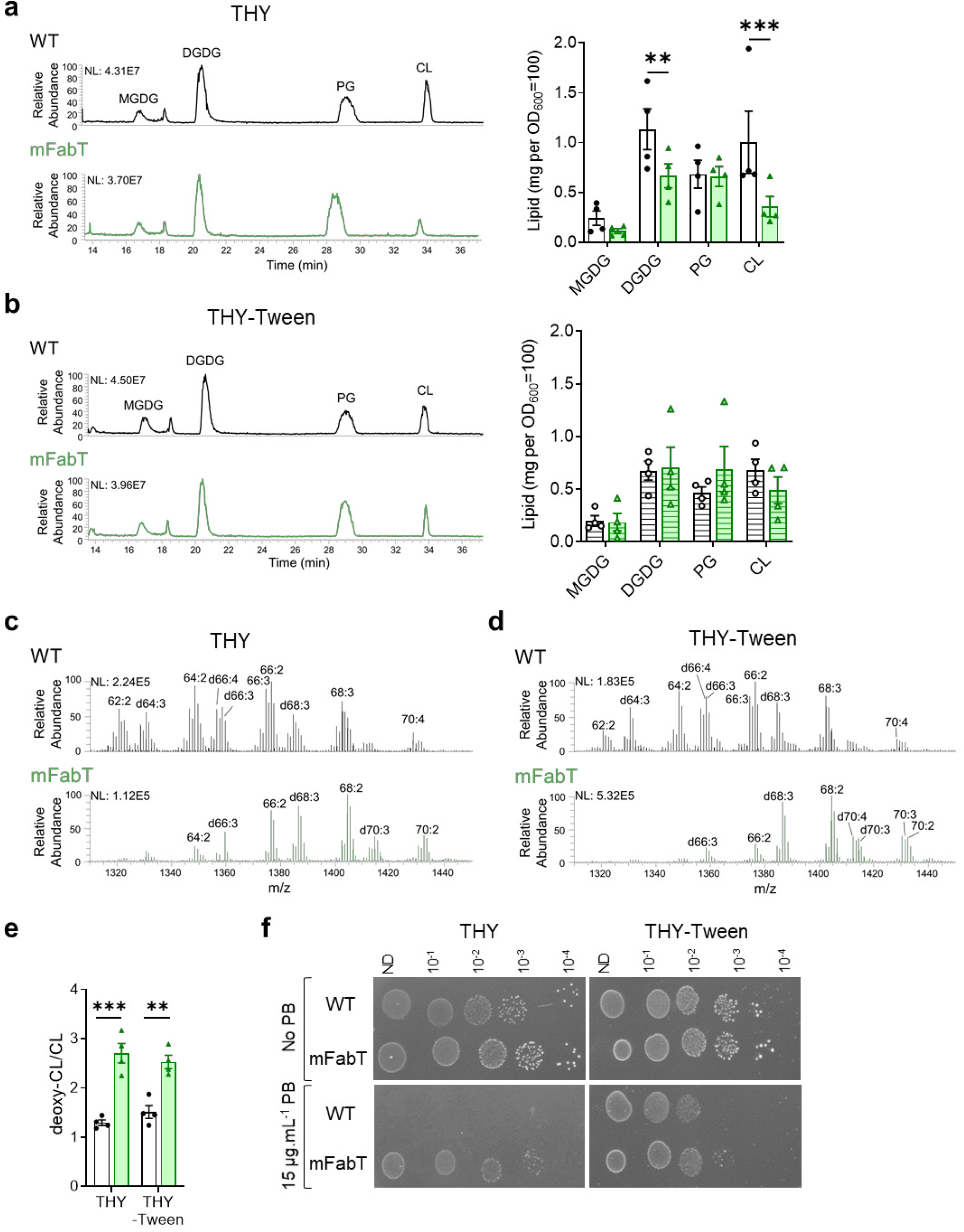
FA differences in WT and mFabT strains impact lipid content and composition. **a, b,** HPLC-MS profiles represent the main lipid classes in WT and mFabT grown in indicated media to OD600=0.5; lipid profiles and quantifications by class. Lipid concentrations correspond to milligrams extracted from OD600=100 equivalents. MGDG, monoglucosyldiacylglycerol; DGDG, diglucosyldiacylglycerol ; PG, phosphatidylglycerol; CL, cardiolipin (Supplementary Table 3). **c, d, e** CL species in WT and mFabT, showing deoxidized cardiolipin (deoxy-CL) species (indicated by a ‘d’ preceding the identification name; see Supplementary Table 2); **a-b,** N=4; **c-d,** N=3. **a, b, e,** 2-way ANOVA, Bonferroni post-test, **p<0.01; ***p<0.001. **f,** Polymyxin B sensitivity. WT and mFabT were precultured to OD600 = 0.5 in THY or THY- Tween and dilutions were spotted on the same respective solid medium supplemented or not with polymyxin B (PB). Plates are representative of 3 independent experiments. **a-d**, NL, normalization level; WT, white bars; mFabT, green bars.

*fabT* mutants are reportedly more resistant to the cationic cyclic peptide polymyxin B ^5,17^. Polymyxin B binds negatively charged lipids such as CL ^18^. Putting together these reports and the CL changes noted above, we examined polymyxin B resistance of mFabT and WT strains in conditions where CL amounts varied (Fig. 1f). Features that decreased CL pools, *i.e.*, *fabT* mutation or FA availability during WT growth, correlated with greater polymyxin B

resistance. These data support the idea that greater polymyxin B resistance in *fabT* mutants is due to lower CL pools, and show that availability of environmental FAs can lead to polymyxin B resistance. They give a rationale for *fabT* mutant emergence upon polymyxin B selection ^17^. However, they do not explain the *in vivo* emergence of mFabT mutants at an early stage of GAS infection.

### Effects of *fabT* on FASII and virulence factor expression

We first examined the basis for the shift towards greater amounts of C18:0 in the mFabT strain when grown in THY medium (Supplementary Table 1). In THY-Tween, FASII genes are repressed in WT, but not detectably in mFabT ^2^. In FA-free THY medium however, the only difference in *S. pyogenes* FASII gene expression was a greater expression of *fabK* in mFabT than in the WT strain, leading to an increased *fabK*: *fabM* ratio in the mFabT mutant, as reported for a *fabT* mutant in *S. pneumoniae* (^2,6^; Supplementary Fig. 4a). FabK and FabM compete for the same substrate to respectively synthesize saturated and unsaturated FAs (Supplementary Fig. 1a); thus, higher *fabK* expression relative to *fabM* is consistent with the higher proportion of saturated FAs in *fabT* mutants ^2,6^. The underlying reason for a higher *fabK*:*fabM* ratio in the mFabT mutant was revealed by analyzing FASII locus organization by qRT-PCR (Supplementary Fig. 4b). All genes from *fabM* to *accD* were cotranscribed. In addition, transcription start sites are present within this operon (Fig. 2a) ^19^. All transcriptional start sites except that of *fabH* are preceded by FabT consensus DNA binding motifs (5’- ANTTTGATTATCAAATT). Notably, two FabT binding sites are predicted upstream of *fabK*, but not of the other FASII transcriptional units (starting positions 400693 and 400931 on the *emm28* M28PF1 genome [CP011535.2] ^13^) ^3^. We suggest that the double FabT binding sites upstream of *fabK*, and basal levels of free FAs in THY are responsible for differential *fabK fabM* regulation and hence increased saturated FA synthesis in the mFabT mutant.

**Fig. 2.**
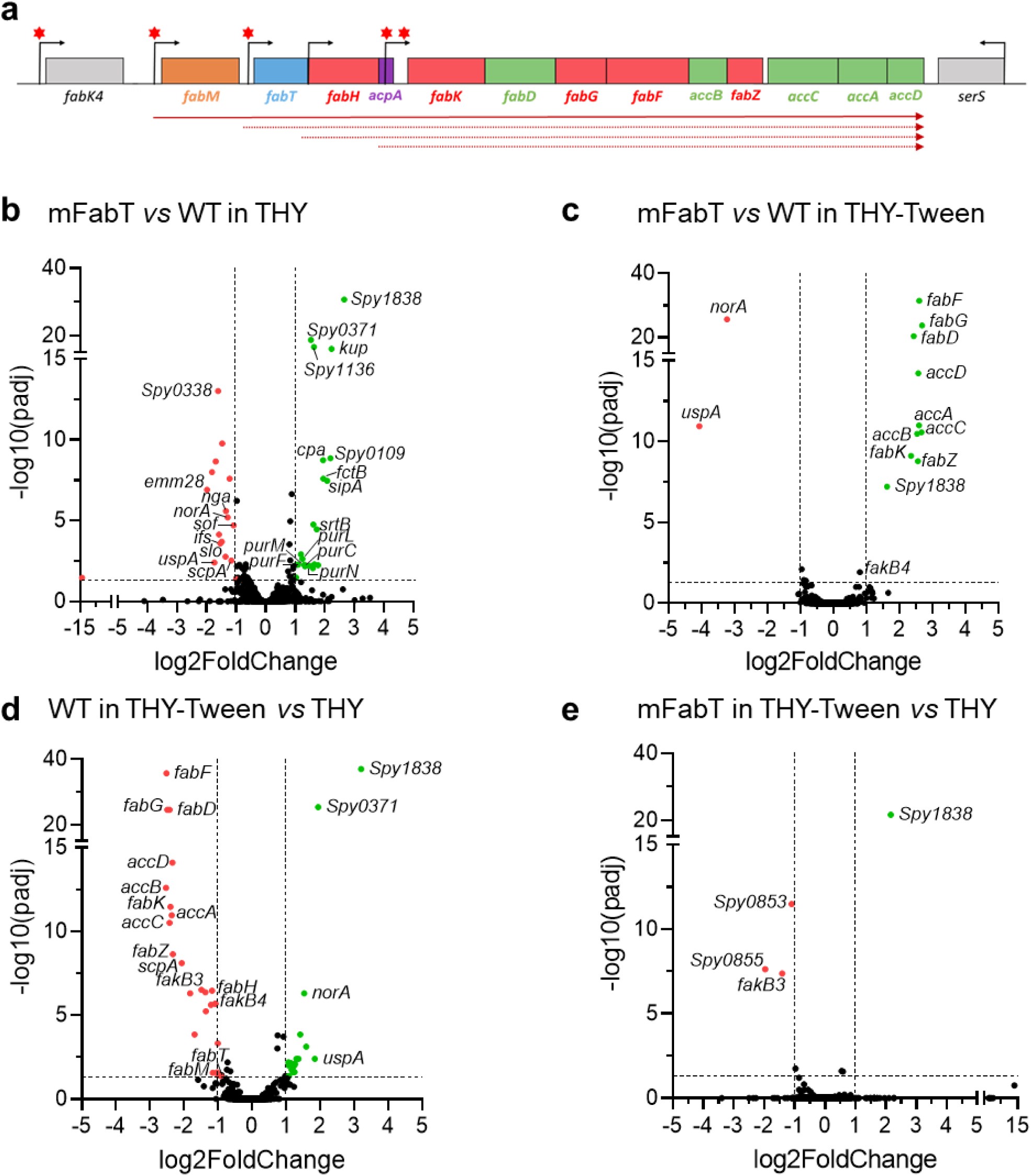
FabT regulon in the presence of eFAs and genetic organization of the GAS FASII genes. **a**, Schematic representation of the GAS FabT regulon, comprising genes of the FASII locus and *fakB4*. Gene positions with names below are represented. Red asterisks, putative FabT binding sites ^3^; bent arrows, transcription start sites; solid arrow, transcript defined by RT-PCRs; dotted arrows, transcripts from previously reported start sites ^19^. **b-e**, Volcano plots of differentially expressed genes, compared as indicated. See supplementary Table 4 for a complete list. Volcano plots were constructed using GraphPad Prism, by plotting the negative base 10 logarithm of the p-value on the y axis, and the log of the fold change (base 2) on the x axis. P-values for comparisons of peak intensities were calculated by t-tests. Padj: calculated p- values were adjusted for multiple testing using the false discovery rate controlling procedure (see Methods section). Gene expression was considered modified in a given condition when the absolute value of log2-Foldchange (FC) was greater than or equal to 1, with an adjusted p-value ≤ 0.05. Data points with low p-values (highly significant) appear at the top of the plot.

We then used RNAseq and qRT-PCR to identify how membrane FA changes might impact and explain the virulence defect during host infection. WT and mFabT expression was compared in THY, and in THY-Tween (as C18:1Δ9 source) to activate WT FabT repression (^3^ for Review; Supplementary Table 4, Fig 2b-e, Supplementary Figure 4c). Several differences, some corresponding to multi-gene operons, distinguished the two strains (Supplementary Table 4). Notably, purine synthesis operon genes (M28_Spy0022 to M28_Spy0026) were upregulated in mFabT, suggestive of increased metabolic activity. An operon encoding adhesins (M28_Spy0107 to M28_Spy0111) was also upregulated; however, expression of virulence genes of the Mga regulon, i.e., C5 peptidase ScpA, and other adhesins, M protein, Sof and SfbX, were down-regulated (Supplementary Table 4). Decreased expression of adhesins and other virulence factors such as SLO (*slo*) and NADase (*nga*) could contribute to the reported poor virulence of *fabT* mutants in infection ^20^.

Addition of C18:1Δ9 activates FabT-mediated repression, which, as reported, turns off FASII gene expression in the WT strain (Fig. 2c, Supplementary Table 4) ^4^. In contrast, FASII genes (see above), and a non-FASII gene within the FabT regulon (M28_Spy1638, encoding a putative fatty acid kinase binding protein; ^14^), remain transcriptionally active in the mFabT mutant, showing that FabT^H105Y^ is defective for FASII repression (Fig. 2e, Supplementary Table 4).

Despite the more pronounced difference in FASII gene expression between WT and mFabT in THY-Tween, only few differences were observed in virulence gene expression. We noted that virulence genes lack FabT binding motifs, suggesting an indirect effect of FabT on their expression. Of note, in THY medium, the mFabT mutation led to longer chain lengths than in WT (determined as C18 to C16 ratios of respectively 2.3 and 1; Supplementary Table 1). In THY-Tween, these ratios are comparable; the WT C18:C16 ratio increased to 2. We speculate that the higher C18:C16 ratio of membrane FAs may provoke lower virulence gene expression, as seen for mFabT in THY, and the WT in THY-Tween (Supplementary Table 4). Thus, the greater proportion of longer FAs in the membrane might be a cause for the decreased expression of virulence genes that lack FabT binding motifs.

### The WT strain has a fitness advantage over mFabT during growth on human decidua

To understand the role of *fabT* and emergence of *fabT* mutants, we designed *ex vivo* assays to study infection with WT and *fabT* variants. mFabT strain capacity to colonize human tissue *ex vivo* was assessed by measuring its growth on human tissue. As *emm28* strains are associated to puerperal fever ^21,22^, we compared *ex vivo* growth of WT and mFabT on human decidua ^23^. In control experiments, both strains failed to grow in RPMI, as assessed by growth ratios between cfus after 8 h incubation and the inocula (Supplementary Fig. 2e). In static conditions, growth of mFabT was 63 % lower than that of the WT strain (Fig. 3a). Growth kinetics of WT*^eryR^*^-i*gfp*^ and mFabT*^eryR^*^-i*gfp*^ GFP-marked strains were then followed in flow conditions at the tissue surface by time-lapse microscopy (Fig. 3b-c). The surface colonized by the WT strain increased throughout the 4 h growth period (Fig. 3b). In contrast, mFabT strain growth only increased during the first half-hour of image acquisition. The average thickness of bacterial microcolonies increased for the WT, but decreased for the mFabT strain (Fig. 3c). Altogether, the WT strain grew with a doubling time of roughly 200 min whereas the mFabT strain did not grow. Thus, in contrast to normal growth in THY medium, the mFabT strain has a major growth defect in the presence of human decidua. These results provide insight into the nature of the colonization defect and virulence attenuation of *fabT* mutant strains ^5,12^.

**Fig. 3.**
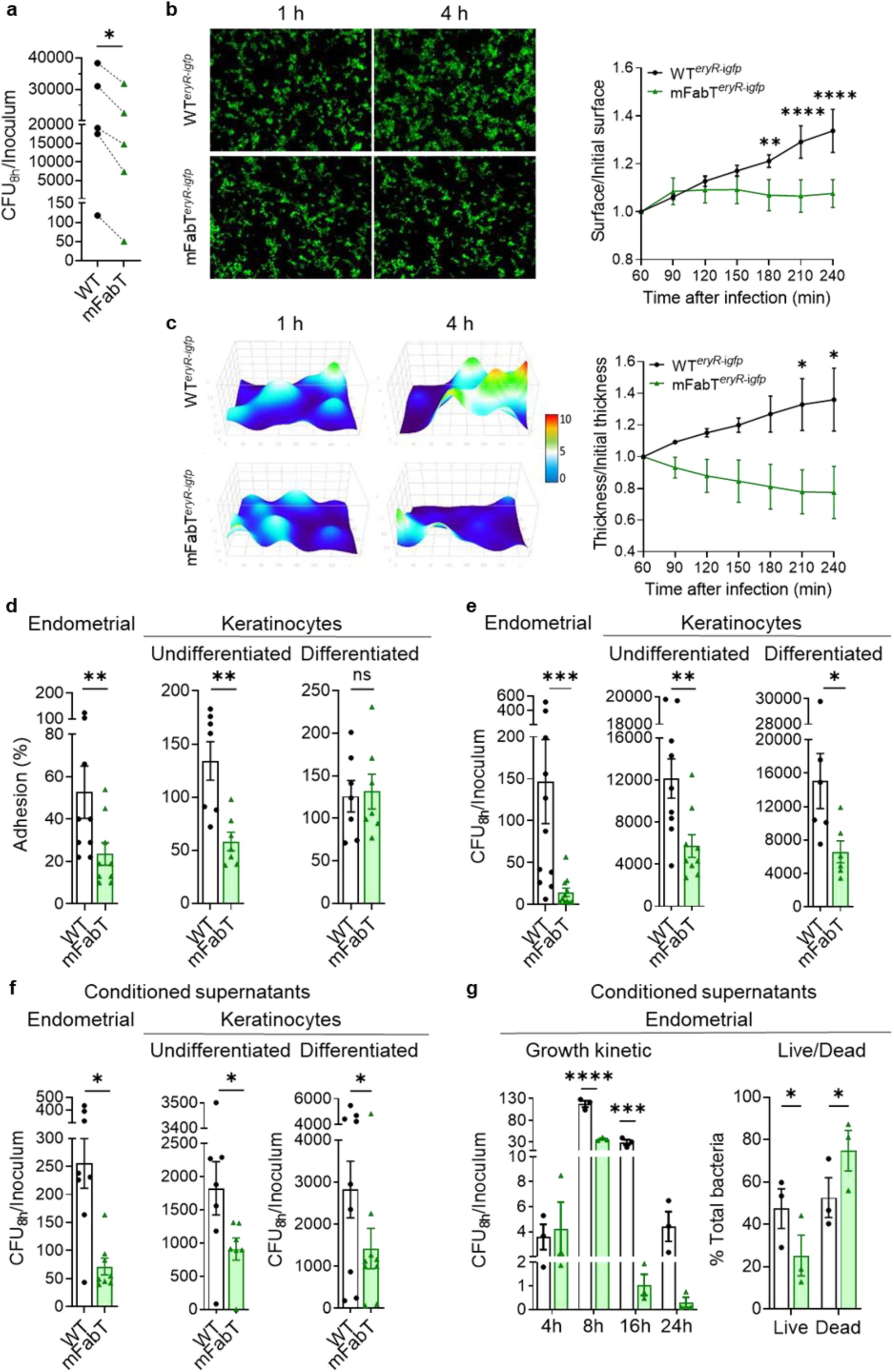
The mFabT strain grows poorly on human tissue *ex vivo* and displays adhesion and growth defects in the presence of human cells or in conditioned cell supernatant. **a**, Comparison of GAS WT and mFabT cfus after 8 h growth in static conditions in the presence of human decidua tissue. **b-c,** Bacterial multiplication at the tissue surface in flow conditions (live imaging); **b, left,** Visualization of WT*^eryR^*^-i*gfp*^ and mFabT*^eryR^*^-i*gfp*^ multiplication in 2D. As poor mFabT growth led to nearly identical images at 1 and 4 h, differences are highlighted by arrowheads; **right,** ratios of areas covered by the two strains. **c, left**, 3D-surface heatmap of bacterial layer thickness at 1 and 4 h. The x, y, and z axes are scaled, color code in µm; **right,** quantification of bacterial layer mean thickness of WT*^eryR^*^-i*gfp*^ and mFabT*^eryR^*^-i*gfp*^ strains on decidual tissue. **d-g**, Comparison of WT and mFabT strain adhesion and growth capacities in the presence of human cells, or conditioned supernatants. Endometrial cells, undifferentiated keratinocytes, and differentiated keratinocytes, and their respective conditioned supernatants were used as specified. **d**, adhesion; **e-f,** growth (cfu.mL^-1^). **g, left,** Bacterial growth kinetics in endometrial conditioned supernatants; **right,** Live/Dead bacteria assessment after 8 h growth in conditioned supernatants. Growth experiments were started with 10^3^ bacteria per ml. Determinations were based on N=5 for **a**, N=3 for **b**, **c**, **g,** N=9, 7, 7 for **d** (left to right), N=11, 9, 6 for **e** (left to right) and N=8, 7, 9 for **f** (left to right). Analyses were done by T test and Wilcoxon test for **a**, **d-f** and 2-way ANOVA, Bonferroni post-test for **b**, **c**, **g**; *p<0.05; **p<0.01; ***p<0.001; ****p<0.0001. **d-f**, Outliers were searched using ROUT method (GraphPad), with Q=1 %. **d-g:** WT, white bars; mFabT, green bars.

### Impaired mFabT fitness is due to defective adhesion and poor growth on human cells and in cell supernatants

Bacterial colonization of host tissue comprises an initial adhesion step, followed by bacterial multiplication ^23^. As GAS has tropism for endometrial and skin tissues, we compared WT and mFabT adhesion capacities on human endometrial cells, and on differentiated (as present in upper skin layers), and undifferentiated skin keratinocytes (Fig. 3d). The mFabT mutant displayed an adhesion defect on endometrial cells as reported ^2^ and undifferentiated keratinocytes compared to the WT. In contrast, the strains adhered similarly on differentiated keratinocytes. The adhesion defect, as also suggested by transcriptome analyses (Supplementary Table 4, Fig. 3d), may contribute to the virulence defect observed in the non-human primate animal models ^5^.

GAS replicates mainly extracellularly during infection of endometrium and skin ^24^. Growth comparisons of WT and mFabT strains on these biotopes, namely endometrial cells, undifferentiated keratinocytes, and differentiated keratinocytes, showed that the mFabT strain was decreased (90 %, 53 % and 56 % respectively) compared to the WT strain (Fig. 3e). The FabT^H105Y^ mutation thus leads to impaired growth in the presence of human cells. To determine whether adhesion is required for the growth differences between WT and *fabT* strains, we assayed bacterial growth in uninfected cell supernatant, termed “conditioned supernatant” (Fig. 3f). The mFabT strain displayed a similar growth defect in endometrial, undifferentiated and differentiated keratinocyte conditioned supernatants (72 %, 50 % and 50 % respectively) compared to the WT. This indicates that cell-secreted products differentially affect growth of WT and mFabT strains. Adhesins differentially produced in WT (Supplementary Table 4, Fig. 3d) could also contribute to promoting higher bacterial densities during infection.

The differences in WT and mFabT growth were visible when grown on endometrial cells or their conditioned supernatants. It is therefore likely that secreted endometrial cell compounds affecting GAS growth are produced independently of infection. Higher bacterial growth densities on differentiated keratinocytes than on conditioned supernatants (compare Fig. 3e and 3f) may be due to greater nutrient availability after infection. Altogether, these data implicate both adhesion and growth defects in poor survival of mFabT in cell infection environments.

Poor mFabT growth suggests that mFabT makes inefficient use of nutrients secreted by eukaryotic cells and/or is more susceptible than WT to secreted bactericidal molecules. We investigated growth kinetics of WT and mFabT strains in endometrial cell conditioned supernatant in time course experiments (Fig. 3g left). Cfu ratios were similar for both strains at 4 h, indicating no difference in the lag time. However, from 8 h onwards, cfu ratios were higher for the WT strain (Fig. 3 f,g left). The use of a live-cell analysis (IncuCyte® Systems, Sartorius, Germany), in which both living and dead bacteria are visualized, revealed that mFabT growth did not peak before the WT, and that both strains grew similarly until ∼3 h; after 3 h mFabT net growth is visibly slower than that of the WT strain (Supplementary Fig. 2f). mFabT mortality was ∼1.5-fold greater than WT at 8 h post-inoculation in conditioned supernatant (Fig. 3g right); these differences are amplified at 16 h and 24 h (Fig. 3g left). We noted that GAS dies rapidly when growth stops, as seen in RPMI (Supplementary Fig. 2e). We conclude that mFabT grows more slowly and dies more rapidly than the WT strain, and lead us to suggest that mFabT death is triggered by poor growth on nutrient-limited medium.

### Faster metabolic turnover in mFabT generates a growth defect during infection

We investigated a possible metabolic basis for the mFabT growth defect, which could reflect an incapacity to use cell-secreted products for growth, and/or higher mortality (Fig. 3g). We used a metabolomics approach to assess metabolites that are differentially consumed by mFabT compared to the WT strain, as performed on conditioned supernatants. Hexoses and amino acids, Asn, Gly, Ile and Lys were the main metabolites overproduced by uninfected cells (Supplementary Fig. 5, Supplementary Table 5). The mFabT strain consumed more hexoses and amino acids Asn, Ile, Lys, and Ser, as seen at 16 h (Fig. 4, Supplementary Table 5). This overconsumption is not linked to a higher bacterial yield, but rather the opposite. It is also not due to faster growth of the mutant relative to WT, but rather to a lower bacterial yield resulting from accrued mortality of this strain (Fig 3g, Supplementary Fig. 2f).

**Fig. 4.**
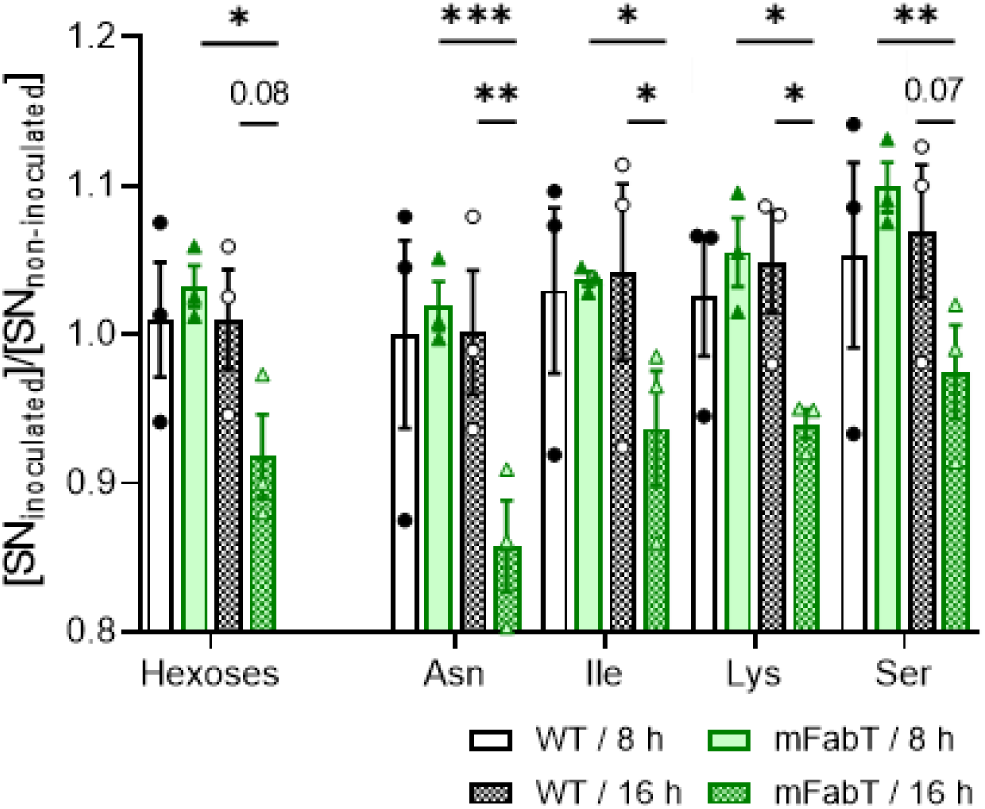
The mFabT strain is more energy-consuming than the WT strain. Metabolomic analysis of conditioned supernatants inoculated with WT or mFabT after 8 or 16 h incubation. Carbohydrate and amino acid consumption by the two strains are compared (see Supplementary Fig. 5 for metabolite levels in noninfected samples, and Supplementary Table 5 for further amino acid results). N=3, 2-way ANOVA, Bonferroni post-test; *p<0.05; **p<0.01; ***p<0.005; p-values just above the p=0.05 threshold are indicated. Strains and growth times are below.

Two characteristics of mFabT may underlie this futile cycle: 1- The high C18:C16 ratio in mFabT compared to WT (respectively 2.3 to 1; Supplementary Table 4) may reduce bacterial fitness by altering activities of membrane components, leading to greater mortality in nutrient-limited medium. 2- Lipid synthesis and FA availability reportedly play a role in regulating metabolic functions ^25^. Continued FASII expression in mFabT may desynchronize coordination between membrane synthesis and growth, such that metabolite import remains active even if other growth factors are unavailable. Consistently, increased mFabT metabolite consumption and greater mortality occurred after 8 h incubation, when growth presumably slows (Fig. 3g, Fig. 4). These data give evidence that the mFabT mutant has greater metabolic consumption at the GAS site of infection, but without growth benefits. Futile energy loss in mFabT might be further due to expression changes associated with its altered membrane FA composition and continued FASII synthesis, leading to increased mortality in nutrient-limited biotopes. These factors might account for the diminished capacity of *fabT* mutants to cause infection ^5^.

### Continued FASII activity in the mFabT mutant leads to lower eFA incorporation

C18:1Δ9 incorporation is reduced in the mFabT strain (Supplementary Table 1). In *S. pneumoniae*, a *fabT* mutant reportedly incorporated only traces of C16:1 ^6^; in an *Enterococcus faecalis fabT* deletion mutant, incorporation of unsaturated FAs, and to a lesser extent, saturated FAs, was defective ^4^. We evaluated FA incorporation by GAS in medium supplemented with C17:1, which is not synthesized by GAS; incorporation of this FA generates a discrete peak by gas chromatography (Fig. 5a left). The proportion of C17:1 was 3-fold higher in the WT than in mFabT (52 %:17 %).

**Fig. 5.**
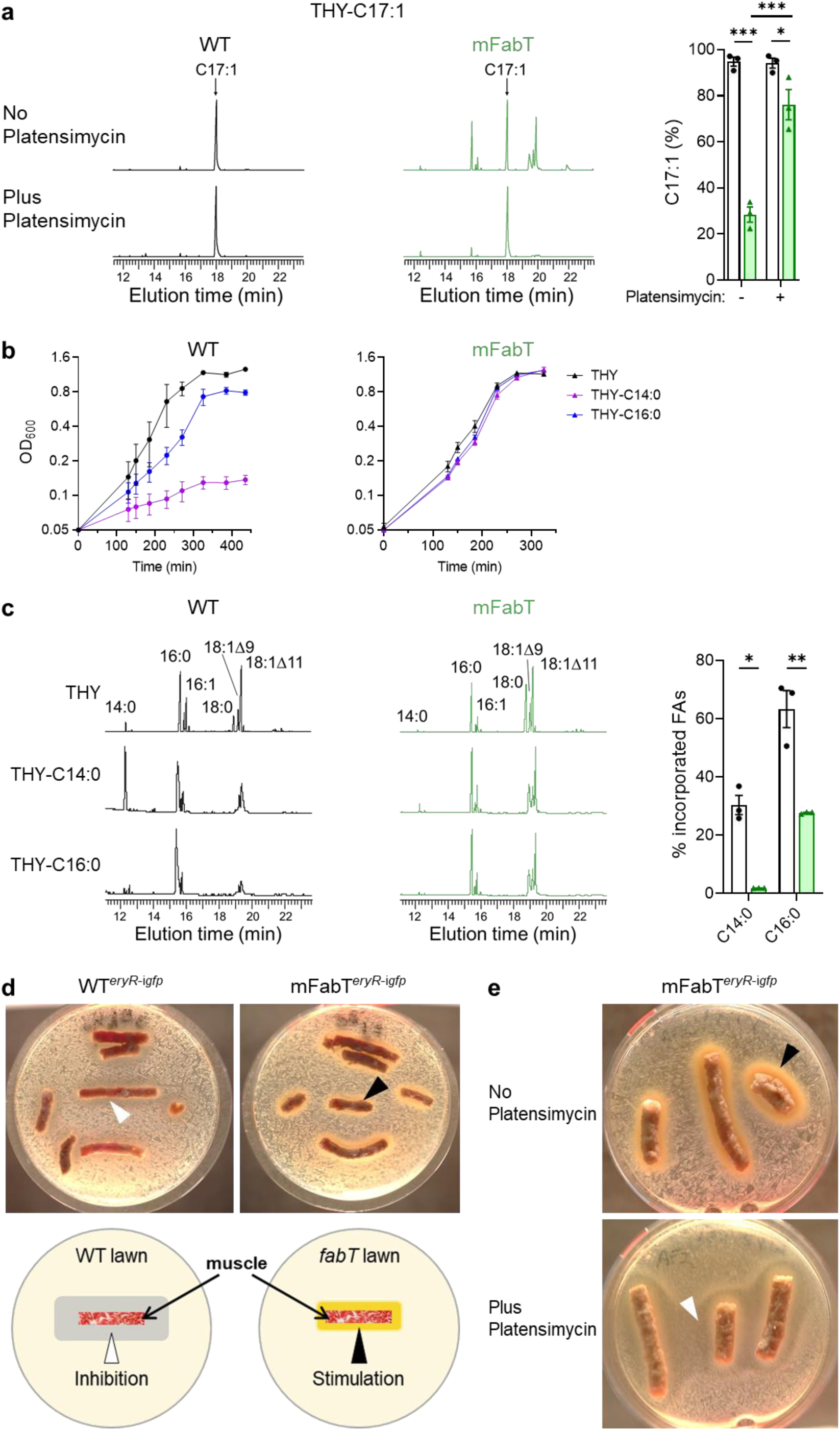
The eFA incorporation defect of mFabT confers a selective advantage in saturated eFA environments, and in a simulated muscle biotope. **a**, FA membrane composition of WT and mFabT strains grown in THY-C17:1 (100 µM), containing or not the FASII-inhibitor platensimycin (1 µg/mL^-1^). **Left**, FA profiles; **right**, quantified proportions of major FAs. Both strains grew in C17:1 regardless of the presence of platensimycin (Supplementary Table 6). **b,** Growth of WT (left), and mFabT (right) in THY supplemented with 100 μM saturated FAs, C14:0 or C16:0 and BSA 0.025%. **c, Left,** Incorporation of exogenous C14:0 and C16:0 from cultures described in ‘**b**’. **Right**, Percent C14:0 and C16:0 incorporation from growth experiments at left. For these analyses, N=3, 2-way ANOVA, Bonferroni post-test, *p<0.05; **p<0.01; ***p<0.001. WT (black lines, white bars) and mFabT (green lines and bars). **d-e,** The indicated strains were spread on solid BHI medium. Pellets of organic ground bovine meat (∼15 % fat), used as muscle source, were placed on the bacterial lawns. Plates were photographed 36 h after incubation at 37 °C. Arrowheads indicate zones of inhibition (white) or growth (black) around muscle sources. N=4. **d, Upper**,WT*^eryR-^*^i*gfp*^ and mFabT^eryR-i*gfp*^ strains were grown in BHI Ery 5, and plated on the same solid medium. **Lower**, schematic representation of GAS strain growth inhibition and stimulation by muscle. **e,** mFabT*^eryR^*^-i*gfp*^ was grown in BHI Ery 5 medium containing C17:1 as FA source, without or with platensimycin to turn off FASII. Cultures were then plated on respectively the same solid medium. N=2.

Continued FASII synthesis in the mFabT mutant might create competition between endogenously synthesized FAs and eFAs for incorporation into membrane phospholipids. To test this, we performed the same experiments in the presence of platensimycin (Fig. 5a right), a FabF inhibitor that blocks FASII synthesis independently of FabT ^26^. WT and mFabT strains grew similarly in the presence of C17:1 and platensimycin (Supplementary Table 6), and C17:1 was the major membrane FA in both strains (Fig. 5a). In conclusion, the FabT^H105Y^ mutant is less responsive to environmental FAs than the WT strain. Poor eFA incorporation in mFabT phospholipids thus correlates with continued expression of FASII genes.

### *De novo* emergence of *fabT* mutants in saturated FA environments

We hypothesized that the defect in eFA incorporation could actually confer a growth advantage to the mFabT strain in toxic lipid environments, and thereby point to conditions of *fabT* mutant emergence. Indeed, FA incorporation can negatively affect bacterial integrity, and free FAs are considered part of the first line of host defense against skin infections ^27^. Notably, increased C18:0 levels in the mFabT mutant compared to WT (^2^; Supplementary Table 1) led us to examine saturated FAs as potential selective pressure for *fabT* emergence. WT and mFabT growth and eFA incorporation were then compared in the presence of C14:0 and C16:0, both of which are found among host lipids ^28^. Both FAs inhibited WT growth, and were nonetheless efficiently incorporated in WT membranes; in striking contrast, mFabT failed to incorporate C14:0, and incorporated >2-fold less C16:0, while strain growth was robust (Fig. 5b-c). This growth advantage incited us to consider that spontaneous *fabT* mutants could emerge in the presence of saturated FAs.

As proof of concept, we grew the WT strain in THY liquid medium without or with C14:0, and then streaked cultures on solid medium containing C14:0. Single colonies appeared on this medium, and *fabT* genes were sequenced. Mutations in *fabT* were obtained in both selection procedures, and encoded FabT variants FabT^T65M^ and FabT^G99S^ (Supplementary Fig. 1b-c). Both these variants were also identified in a primate infection study ^5^. We compared growth and FA-incorporation phenotypes of the isolated FabT^T65M^ and FabT^G99S^ mutants, as well as Δ*fabT* mutant as reference, to those of WT and mFabT strains Supplementary Fig. 6. Growth at 4h of the newly isolated mutants, mFabT and Δ*fabT*, was essentially unaffected by C14:0. Moreover, the mutant expressing FabT^T65M^, mFabT and Δ*fabT* strains incorporated ≤ 2 % C14:0 compared to ∼53 % ± 6 in the WT. The mutant expressing FabT^G99S^ incorporated 20 % ± 3, well below that in the WT strain. These results provide a rationale for emergence of *fabT* mutants, by generating a growth advantage in lipid-containing biotopes as may be present at the infection locus.

### Evidence that eFA exclusion by mFabT confers a growth advantage in a simulated muscle biotope

We hypothesized that *fabT* mutations could provide a transient advantage in lipid-rich biotopes, which would explain their emergence in muscle ^5^. An *ex vivo* model was devised to assess effects of muscle on GAS growth. For this, commercial organic 15 % fat meat, *i.e.*, muscle from cattle, was placed on lawns of WT*^eryR^*^-i*gfp*^ and mFabT*^eryR^*^-i*gfp*^. This is within the percent fat range estimated for human muscle (8-30 %; in contrast, uterine fluid lipid concentrations are >1000-fold lower ^29,30^). Remarkably, WT strains developed a weak inhibitory halo surrounding the muscle samples. In strong contrast, the mFabT strain grew directly and vigorously around muscle sources (Fig. 5d).

This result is consistent with growth inhibition of WT but not mFabT GAS, notably by saturated eFAs (Fig. 5b). To confirm that vigorous growth of mFabT around muscle relates to its lower incorporation of FAs from muscle lipids, we tested the effects of adding the FASII inhibitor platensimycin to growth medium and plates. All media contained C17:1, which is incorporated and allows mFabT growth in the presence of the FASII inhibitor (Fig. 5a, Supplementary Table 6). As in Fig. 5d, mFabT growth was robust around muscle sources in the absence of FASII inhibitor (Fig. 5e upper). In contrast, growth was strongly inhibited around the muscle source when platensimycin was present (Fig. 5e lower). The marked inhibitory effect of the muscle sample on the *fabT* mutant when FASII is blocked gives strong evidence that incorporation of muscle lipids negatively affects GAS growth. We note that indirect effects, consequent to FASII inhibition by platensimycin and/or by FabT repression, may contribute to growth inhibition. We conclude that continued FASII activity in mFabT mutants reduces eFA incorporation and thus protects bacteria from toxicity of host lipids as present in muscle biotopes.

## Discussion

Our work establishes the causal origin for *fabT* mutation emergence and provides an explanation for its disappearance during invasion (Fig. 6): GAS is genetically designed to incorporate eFA from lipid-containing environments, and repress FASII. We showed that WT GAS growth is inhibited by saturated FAs in such environments. Counter-selection can lead to emergence and outgrowth of *fabT* mutants, which explains their detection at a non-invasive step of infection ^5^. At this step, the *fabT* mutation would confer a transient advantage over non-mutant strains. We first showed the *fabT* growth advantage, and mutant emergence, using C14:0 selection. We then demonstrated that the lipid-rich inter-muscular environment allows *fabT* mutant growth in a FASII-dependent manner, while inhibiting the WT strain. While the overall muscle and uterine fluids environments where GAS multiply are very different, it is notable that their lipid concentrations, differ by ∼1000-fold (150 mg/g muscle, compared to 20-200 µg/ml in uterine fluid ^29,30^). We suggest that exposure to high inter-muscular lipid availability would contribute to WT mortality. However, the *fabT* mutation has a cost for virulence: it results in a multiplication defect and higher mortality in the presence of human cells, which is confirmed on human decidua tissue and its fluids. Continued FASII activity in *fabT* mutant strains provokes a state of futile bacterial metabolism where increased metabolite uptake does not lead to improved growth.

**Fig. 6.**
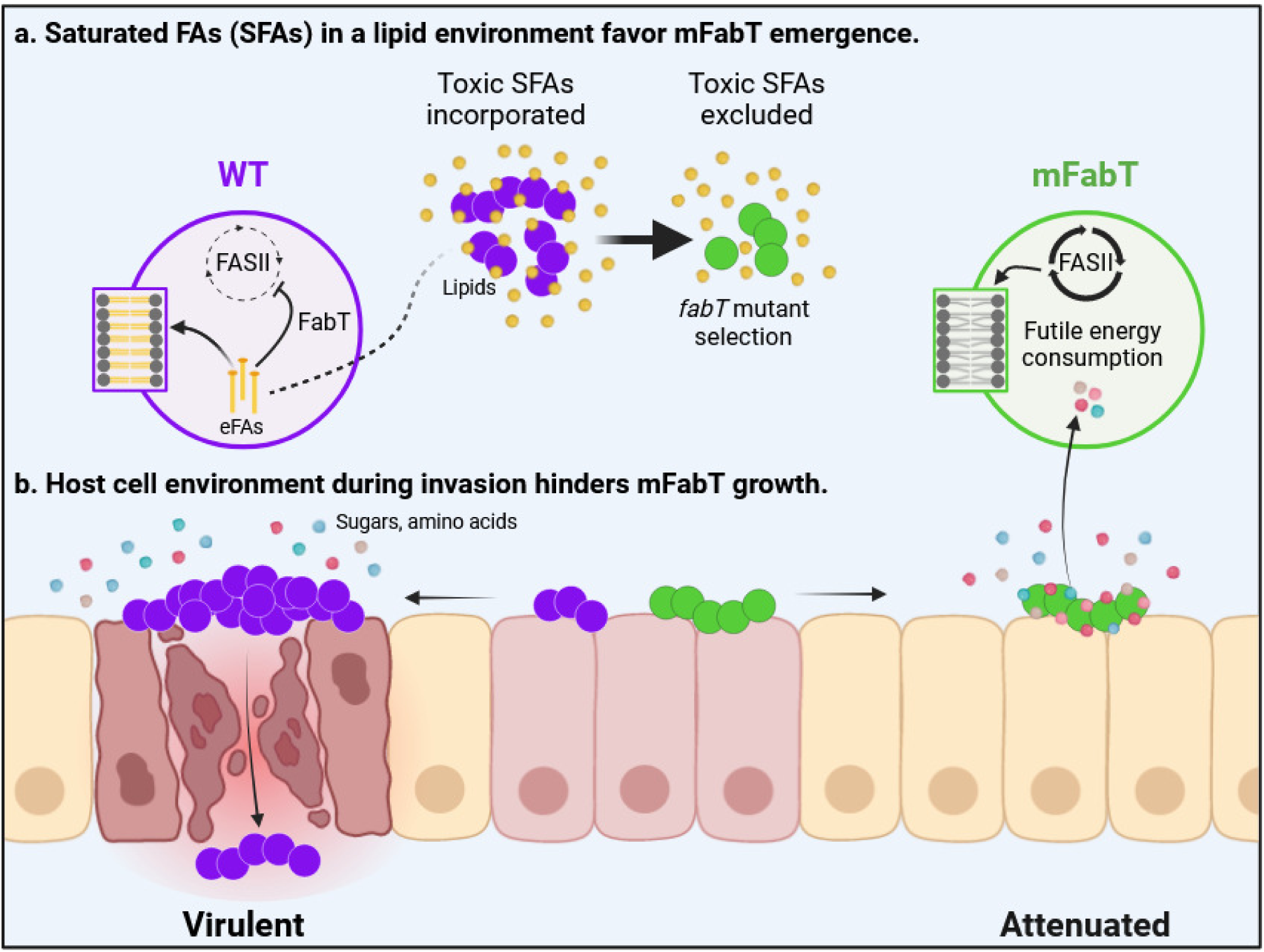
Model for emergence of *fabT* mutants that are attenuated for virulence. a, Saturated FAs (SFAs) in a lipid environment favor mFabT emergence. Toxic FAs may be present in initial GAS contacts with the host. Counter-selection would lead to emergence of FA-insensitive *fabT* mutants, conferring a growth advantage. In a proof of concept, we show that *fabT* mutants are selected in an SFA environment. **b, Host cell environment during invasion hinders mFabT growth**. Compared to the WT, *fabT* mutant bacteria fail to develop and die more rapidly when exposed to human cells; they are also impaired for adhesion. Longer SFAs and continued FASII activity in *fabT* mutants provokes a state of futile bacterial metabolism where metabolite uptake is stimulated, but does not lead to improved growth. Thus, *fabT* mutants in GAS populations may confer a survival advantage at the inoculation site, but do not withstand host cell infection conditions. Mauve and green circles, WT and mFabT cocci; zoom is on phospholipids. Small yellow circles and lines, lipids and eFA hydrolysis products respectively, small red, blue, pink circles, sugars and amino acid residues. Figure was drawn using BioRender (BioRender.com).

Our findings indicate that futile FASII synthesis by mFabT is detrimental for GAS virulence. We showed previously that blocking FASII with antibiotic inhibitors, mutation, or deletion did not prevent infection by *Streptococcus agalactiae*, nor by other Firmicute pathogens ^31–33^. FASII inhibition and eFA incorporation in phospholipids corresponds to the natural feedback inhibition in response to eFAs ^3,31^. The contrary, *i.e.*, making FASII synthesis constitutive by inhibiting FabT, is detrimental for *in vivo* infection. FabT is thus a promising target for new therapeutics against specific Gram-positive pathogens including GAS, *S. agalactiae*, *S. pneumoniae*, and *E. faecalis*.

## Methods

### Bacterial strains and culture conditions

The strains used in this study are described in Supplementary Table 7. GAS strains were grown under static condition at 37 °C in Todd Hewitt broth supplemented with 0.2 % Yeast Extract (THY) or on THY agar (THYA) plates, or in brain heart infusion (BHI) liquid or agar medium when specified. Medium was supplemented with 0.1 % Tween 80 (THY-Tween; Sigma-Aldrich, Ref. P1754) as indicated, as a source of C18:1Δ9. THY was also supplemented with the saturated FAs, C14:0 and C16:0, or the unsaturated FA C17:1 (Larodan, Sweden) in the presence of FA-free bovine serum albumin (0.025 %, Sigma-Aldrich, Ref. A6003) The FASII inhibitor platensimycin (Tebubio, France) was added at (0.5 to 1 µg/ml) as indicated. For WT*^eryR^*^-i*gfp*^ and mFabT*^eryR^*^-i*gfp*^ (Supplementary Table 7) strains, medium was supplemented with 5 to 10 μg/ml of erythromycin (Ery). Strains were prepared as follows, unless specified otherwise: overnight cultures were diluted to an OD600 = 0.05 and grown in THY to the exponential phase (OD600 comprised between 0.4 and 0.5). For GFP expression, exponential-phase bacteria were further diluted to OD600 = 0.1 in THY supplemented with 10 μg/ml erythromycin, and 20 ng/ml anhydrotetracycline to induce GFP expression, grown for 90 min at 37 °C, and diluted in RPMI as indicated below. For growth in saturated FAs, WT and mFabT THY precultures were diluted in THY or THY-C14:0 or THY-C16:0 to OD600 = 0.05, transferred to 50 mL falcon tubes, and incubated at 37 °C. Growth was determined by OD600 readings at designated time points.

### Strain constructions

Primers used for cloning and strain verification are described in Supplementary Table 8. The Δ*fabT* strain corresponds to a *fabT* deleted mutant. It was obtained by homologous recombination of the plasmid pG1-Δ*fabT* following a published protocol ^34,35^. The DNA fragments encompassing *fabT* were cloned in BamHI – EcoRI digested pG1 using the In Fusion cloning kit® (Clonetech). This led to the deletion of *fabT* from nucleotides 49 to 389, as confirmed by PCR. The mFabT*^eryR^*^-i*gfp*^ strain harboring an integrated inducible *gfp* gene was constructed as described for the WT*^eryR^*^-i*gfp*^ strain ^23^. Whole genome sequencing was performed on the mFabT and mFabT*^eryR^*^-i*gfp*^ constructed strains, and no surreptitious mutations were found (Bioproject PRJNA926803, accession number SAMN34247893 for mFabT, SAMN34247911 for mFabT*^eryR^*^-i*gfp*^).

### Fatty acid analysis

Strains were grown until OD600 = 0.4 - 0.5. Fatty acids were extracted and analyzed as described ^2,14,31–33^. Briefly, analyses were performed in a split-splitless injection mode on an AutoSystem XL Gas Chromatograph (Perkin-Elmer) equipped with a ZB-Wax capillary column (30 m x 0.25 mm x 0.25 mm; Phenomenex, France). Data were recorded and analyzed by TotalChrom Workstation (Perkin-Elmer). FA peaks were detected between 12 and 40 min of elution, and identified by comparing to retention times of purified esterified FA standards (Mixture ME100, Larodan, Sweden). Results are shown as percent of the specific FA compared to total peak areas (TotalChrom Workstation; Perkin Elmer).

### Lipid analysis

Strains were grown as 200 ml cultures in THY or THY-Tween until OD600 = 0.4 - 0.5. Lipid extractions and identifications were performed as described ^32,36–38^. Lipid separation was realized by normal phase HPLC (U3000 ThermoFisher Scientific) using a Inertsil Si 5µm column (150 x 2.1 mm I.D.) from GL Sciences Inc (Tokyo, Japan). Lipids were quantified using a Corona-CAD Ultra and identified by mass-spectrometry negative ionization and MS^2^/MS^3^ fragmentations (LTQ-Orbitrap Velos Pro). The adducts observed for molecular ions were: CH3COO- for MGDG, CH3COO- and H- for DGDG, H- for PG and CL. The concentration of each lipid class was determined as described using as standards (from Avanti, Germany) DGDG, 840524P-5MG; MGDG, 840523P-5MG; CL (heart CA), 840012P-25MG; PG (egg), 841138P-25MG ^39^. Lipid spectra were analyzed on Xcalibur^TM^ software (ThermoFisher Scientific, version 4.2.47). Lipid concentrations are presented as milligrams per OD600 = 100 for all samples. The Lipid Maps ‘Structure search’ program https://www.lipidmaps.org/resources/tools/bulk-structure-search/create?database=COMP_DB was used to identify FA moieties on identified lipid species.

### Polymyxin B assay

Polymyxin B sensitivity was assayed as described ^5^. Bacteria were grown to OD600 = 0.4 - 0.5 in THY or THY-Tween. Serial dilutions were prepared in PBS, and 2.5 µl of each dilution was inoculated onto THY or THY-Tween plates containing or not 20 µg.ml^-1^ polymyxin B (Sigma-Aldrich, Ref. 81271). Plates were incubated at 37 °C for approximately 24 h and photographed. Experiments were done in biological triplicates.

### In silico analysis

Geneious prime Biomatters development, www.geneious.com was used to identify 5’-ANTTTGATTATCAAATT-3’, the putative FabT binding sequence, on the M28PF1 genome, accepting up to 2 mismatches.

### RNA isolation and Illumina RNA-seq sequencing

GAS strains were cultured at 37 °C in THY or THY-Tween, and cells were harvested during exponential growth (OD600 between 0.4 and 0.5). Independent triplicate cultures were prepared for each condition. For RNA preparation, 2 volumes of RNA protect* (Qiagen) was added to cultures prior centrifugation (10 min 12,000 g) and total RNA was extracted after lysing bacteria by a 30 min 15 mg.ml^-1^ lysozyme, 300 U.ml^-1^ mutanolysin treatment at 20 °C followed by two cycles of Fast-prep (power 6, 30 s) at 4 °C. RNA extraction (Macherey-Nagel RNA extraction kit; Germany) was done according to supplier instructions. RNA integrity was analyzed using an Agilent Bioanalyzer (Agilent Biotechnologies, Ca., USA). 23S and 16S rRNA were depleted from the samples using the MICROBExpress Bacterial mRNA enrichment kit (Invitrogen, France); depletion was controlled on Agilent Bioanalyzer (Agilent Biotechnologies). Libraries were prepared using an Illumina TS kit. Libraries were sequenced generating 10,000,000 to 20,000,000 75-bp-long reads per sample.

### RNA-Seq data analysis

The MGAS6180 strain sequence (NCBI), which is nearly identical to M28PF1 ^13,21^, was used as a reference sequence to map sequencing reads using the STAR software (2.5.2b) BIOCONDA (Anaconda Inc). RNA-seq data were analyzed using the *hclust* function and a principal component analysis in R 3.5.1 (version 2018-07-02). For differential expression analysis, normalization and statistical analyses were performed using the SARTools package and DESeq2 ^40,41^ *p*-values were calculated and adjusted for multiple testing using the false discovery rate controlling procedure ^42^. We used UpsetR to visualize set intersections in a matrix layout comprising the mFabT *versus* the WT strain grown in THY and in THY-Tween, and growth in THY-Tween *versus* THY for each strain ^43,44^. Transcriptomic data was deposited in the Mendeley database https://data.mendeley.com/datasets/68bhhsy2p4/1.

### *Ex vivo* GAS growth capacity analysis

Human placentas with attached maternal-fetal membranes were collected and processed as described ^23^ with the following modifications. Tissues were obtained after vaginal delivery, and the samples used were from regions located far from the cervix, known as zone of intact morphology ^45^. Human decidual explants were infected within hours of their reception.

*Ex vivo* bacterial growth capacity in the presence of decidua human tissue was done as follows: exponentially growing GFP-expressing bacteria were washed twice in PBS and diluted in RPMI at a final concentration of 10^4^ bacteria per ml. Decidua tissue were washed twice in PBS. One ml of bacteria were then added to tissue, followed by incubation at 37 °C + 5 % CO2. After 8 h, the tissues were shredded using Precellys Evolution (Bertin Technologies) (6 x 20 s at 5500 rpm with a 20 s pause between shaking). Serial dilutions of shredded material were plated on THYA plates. The number of cfus was determined after 24 h of growth at 37 °C and normalized to the inoculum for each experiment.

Live bacterial multiplication on human tissue: infection of maternal-fetal explants, image acquisition and treatments were realized as described ^23^.

### Study approval

The study of the human maternal-fetal membranes was approved by the local ethics committee (Comité de Protection des Personnes Ile de France III, no. Am5724-1-COL2991, 05/02/2013). All participants provided written informed consent prior to inclusion in the study at the Department of Obstetrics, Port Royal Maternity, Cochin University Hospital, Paris, France.

### Cell culture

HEC-1-A (ATCC_ HTB-112TM) endometrial epithelial cells were cultured as described ^46^, in McCoy’s 5A medium (Gibco, Ref. 26600080) supplemented with 10 % fetal bovine serum at 37 °C, 5 % of CO2. HaCaT (Addex-Bio T0020001) keratinocytes were cultivated as recommended, in DMEM high glucose medium (Gibco, Ref. 31966) supplemented with 10 % fetal bovine serum at 37 °C, 5 % of CO2. HaCaT cells were maintained in the undifferentiated state by cultivation in poor DMEM medium (Gibco, Ref. 21068-028) supplemented with 1X glutaMax (Gibco, Ref. 35050-038), 1X sodium pyruvate (Gibco, Ref. 11360-039), 2 % fetal bovine serum and 8 % chelated fetal bovine serum using Chelex^®^ 100 Resin (BioRad, Ref. 142- 1253). To differentiate HaCaT cells, 2.8 mM CaCl2 (Sigma-Aldrich, Ref. 21115) was added to the medium. Cells were differentiated after seven days as described ^46^. All cell lines were routinely evaluated for cellular morphology and growth characteristics by microscopy.

### Bacterial adhesion capacity

GAS adhesion capacity was evaluated as described after growing bacteria in THY to an OD600 of 0.4 to 0.5 ^34^. Values were normalized to the inoculum for each experiment.

### Bacterial growth capacity in the presence of eukaryotic cells or culture supernatants

GAS were cultured in THY to OD600 = 0.4 - 0.5. Bacteria were washed twice in PBS and diluted in RPMI medium without glutamine (Gibco, Ref. 32404-014) to infect cell cultures or inoculate filtered cell culture supernatants (conditioned supernatants) at a final concentration of 10^3^ and 10^4^ bacteria per ml, respectively. Confluent cells in 24-well plates were starved 24 h before the experiment, *i.e.* incubated in RPMI medium without glutamine, and washed twice in PBS. Cells were infected with 1 ml of bacteria and incubated at 37 °C + 5 % CO2. After 8 h, supernatants were recovered and cells were lysed with 1 ml distilled water. The fractions were pooled and serial dilutions plated on THYA plates. Conditioned supernatants were prepared by incubating cells in 1 ml RPMI at 37 °C + 5 % CO2 for 8 h. The conditioned supernatant was recovered, inoculated with 10^3^ bacteria, and incubated for another 4, 8, 16 or 24 h. Serial dilutions were plated on THYA plates. The number of cfus was determined after 24 h of growth at 37 °C and normalized to the inoculum for each experiment.

### Live / dead analysis

After bacterial growth in the HEC-1-A conditioned supernatant during 8 h, bacterial mortality was determined using the LIVE/DEAD® BacLightTM Bacterial Viability Kit (ThermoFisher Scientific, Ref. L7012) as described for flow cytometry utilization using an ACCURI C6 cytometer (BD Biosciences, Le pont de Claix,France) from the CYBIO Core Facility. Bacteria were grown in HEC-1-A conditioned supernatant for 8 h for testing. Results of three independent experiments were analyzed using the BD Accuri C6 software.

### Metabolomic analysis

HEC-1-A conditioned supernatants were inoculated or not with WT or mFabT strains during 8 or 16 h and prepared as described above (see ‘Bacterial growth capacity in the presence of eukaryotic cells or culture supernatants’). The metabolite composition of these supernatants was analyzed by Proteigene (https://proteigene.com) using MxP® Quant 500 kit (Biocrates) by two analytical methods, LC-MS/MS for small molecules and FIA-MS/MS for lipids. This analysis was repeated on 3 independent series of supernatants and on RPMI. These analyses have a defined detection threshold (LOD) for each family of metabolite. Metabolomic data were deposited in the Mendeley database: https://data.mendeley.com/datasets/5cnyvy7vvf/2.

### Spontaneous *fabT* mutant isolation

WT strain overnight precultures were diluted in THY and grown at 37 °C. When the THY culture reached mid-exponential phase, (OD600 = 0.4 - 0.5), bacteria were either diluted 1/10 and transferred to THY-C14:0 supplemented with BSA 0.025 %, or 1.5x10^7^ cfu were streaked on THYA supplemented with C14:0 (THYA-C14:0). The THY-C14:0 liquid culture and plates were incubated 60 h at 37 °C. THY-C14:0 liquid cultures were subsequently streaked on THYA-C14:0 plates. Thirteen and six colonies were isolated, respectively from the THY-C14:0 liquid then THYA-C14:0 plates, and THYA-C14:0 plates. Isolated colonies were subsequently grown on THYA. Eight colonies issued from the liquid selection, and all six clones from the solid medium were used for *fabT* sequencing; PCR was performed directly on patched colonies. The oligonucleotides used were FabT-222, and FabTavComp, (Supplementary Table 8) binding 221 bp 5’ from the T of the TTG translation start site and 565 bp downstream of it, respectively. The *fabT* gene and surrounding sequences were amplified by PCR using the Green Taq DNA Polymerase, GenScript, according to the manufacturer’s instruction, with 30 cycles with a hybridizing temperature of 50 °C and an elongation time of 1 min. Sanger sequencing was carried out by Eurofin Genomics (https://eurofinsgenomics.eu/en/custom-dna-sequencing/portfolio-overview/) on PCR products. These analyses identifed FabT^T65M^ in all eight sequenced clones from liquid selection, and FabT^G99S^ in one of six clones on solid medium; the other clones displayed a WT FabT.

### *Ex vivo* assessment of GAS WT and mFabT growth on muscle (meat) lipids

WT*^eryR^*^-i*gfp*^ and mFabT*^eryR^*^-i*gfp*^ strains (Supplementary Table 7) ^23^ were grown overnight in BHI (Ery 5), and then diluted to OD600 = 0.05 in FA-free bovine serum albumin (referred to as BSA; 0.025%) plus C17:1 100 µM in 1 ml, without or with platensimycin 1 µg/ml, with Ery 5 for selection. After 4 h growth, culture densities were adjusted to OD600 = 1 in BHI, and 35 µl were spread on 5 cm diameter plates containing 5 ml BHI agar plus BSA 0.025 % and C17:1 100 µM, without or with platensimycin 0.5 µg/ml, with Ery 5 for selection. The muscle samples used were from organic cattle meat bought frozen and pre-ground, containing 15 % fat (as declared by supplier; Picard, France). Absence of contaminants was checked by plating without bacteria. Samples were placed directly on the lawns, and plates were incubated 48 h at 37 °C, and photographed. N=4 for experiments without platensimycin, and N=2 for those with platensimycin.

## Statistical analysis

Data were analyzed with GraphPad Prism version 9.4.1. The tests used are indicated in figure legends. Statistical significance is indicated by: ns (not significant, p > 0.05); *, p < 0.05; **, p < 0.01; ***, p < 0.001; ****, p < 0.0001. In addition, the Shapiro-Wilk normality test was used for all t-tests.

## Data availability

All relevant data are within the manuscript, supporting information files, and depositories. Transcriptome, metabolome data were deposited in the Mendeley repository (respectively https://data.mendeley.com/datasets/68bhhsy2p4/1 and https://data.mendeley.com/datasets/5cnyvy7vvf/2). Lipidomics were deposited in the Mendeley database in MZML open-source format (https://data.mendeley.com/datasets/cf578v8d8b/1). All data are publicly available as of publication. Any additional information required to reanalyze the data reported in this paper is available from the authors upon request.

## Supplementary Figures

**Supplementary Fig. 1 | FabT regulator and FASII pathway in GAS. a,** The FASII synthesis pathway comprises a first initiation phase for precursor synthesis, followed by the recursive elongation cycle. The final product, acyl-ACP, supplies FAs for phospholipid synthesis. FabM (orange) leads to unsaturated *cis* FAs; FabK products are saturated. Initiation phase and elongation cycle enzymes are represented in green and red, respectively. From ^1^. **b,** FabT sequence; amino acids involved in DNA binding are in red, and those interacting with acyl-Acyl carrier protein are in blue. Arrow indicates the His105Tyr FabT mutation studied in this work. Magenta stars highlight amino acids spontaneously mutated *in vivo* and in a saturated- FA environment (this work). **c,** Overall structure of FabT dimer predicted by Alphafold and adapted with ChimeraX (see references 1-2 in supplementary Methods); one monomer is represented as multicolored (each color designates a separate domain), and the other is beige. Residues Thr65, Gly99 and His105, in magenta, correspond to mutants isolated in this study.

1. Lambert, C., Poyart, C., Gruss, A. & Fouet, A. FabT, a Bacterial Transcriptional Repressor That Limits Futile Fatty Acid Biosynthesis. Microbiol Mol Biol Rev, e0002922, doi:10.1128/mmbr.00029-22 (2022).

**Supplementary Fig. 2 | Impact of FabT mutations on GAS growth. a-d,** WT growth was compared to mFabT (**a** and **b** left; adapted from reference 1) or to Δ*fabT* (**c** and **d**) on THY and THY-Tween. Viability tests (**a** and **b,** right), using the LIVE/DEAD® BacLightTM Bacterial Viability Kit, were performed on WT and mFabT cultures after growth to OD600 = 0.4 - 0.5. Legend at the right of **d** is for growth curves. **e,** Ratios of CFUs of WT or mFabT strains after 8 h over respective initial inocula (10^3^/ml for each) in RPMI medium; ratio below 1 indicates that bacteria die. **f**, Real-time growth of WT and mFabT in endometrial conditioned supernatant followed using Live-Cell Analysis System (IncuCyte®, Sartorius) and the Basic analyzer. **a-d**, **e**, N=3; **f** N=10; differences in **a, b,** and **e** were not statistically significant using T-test. **a, b, e**, Outliers were searched using ROUT method (GraphPad), with Q=1 %. WT, black lines or white bars; mFabT, green lines or bars.

1.Lambert, C. et al. Acyl-AcpB, a FabT corepressor in Streptococcus pyogenes. J Bacteriol 205, e0027423, doi:10.1128/jb.00274-23 (2023).

**Supplementary Fig. 3 | Phospholipid membrane composition.** Identification of **a**, monoglucosyldiacylglycerol (MGDG), **b**, diglucosyldiacylglycerol (DGDG), **c**, phosphatidylglycerol (PG), **d**, cardiolipin (CL) and deoxidized cardiolipin (Deoxy-CL). For each class, lipids are presented as the percentage of total lipids, and are quantified in Supplementary Table 2. For **a**, **b**, and **c**, fatty acid assignments are presented directly on mass spectrometry analyses. Cardiolipin assignments were ambiguous and are not shown. Statistical values were determined using 2-way ANOVA, Bonferroni post-test. *p<0.05; **p<0.01;***p<0.001; ****p<0.0001. Strains were grown in THY (open bars) and THY-Tween (hatched bars). WT, black lines and white bars; mFabT, green lines and bars.

**Supplementary Fig. 4 | Impact of mFabT on FASII and non-FASII gene expression in the absence of FA supplementation**. **a-c**, WT and mFabT strains were grown in THY medium. **a, c**, RNAs were quantified by qRT-PCR. Expression was normalized to that of *gyrA;* relative gene expression is expressed as the log2-fold ratio. **a,** FASII gene expression of mFabT relative to WT is reproduced from Reference 1**. b**, mRNA transcript analysis of FASII locus genes: agarose gel of PCR amplification products on cDNA using primer pairs from neighboring genes. MW, molecular weight reference (Generuler 100 bp, ThermoFisher Scientific). Lane 1, *fabM*-*fabT* (234 bp); 2, *fabH*-*acpA* (221 bp); 3, *acpA*-*fabK* (302 bp); 4, *fabZ*-*accC* (105 bp); 5, *accD*-*serS* (288 bp). Results confirm the FASII transcriptional units indicated in Fig. 2a. **c**, expression of non-FASII genes in WT and mFabT strains (see Supplementary Table 1 for gene assignments). The relative gene expression is expressed as the log2-fold ratio in a given strain grown in THY-Tween vs in THY. *, significance of differences in the two media between WT and mFabT. In **a**-**c,** WT, white bars; mFabT, green bars. **a, b**, N=4; **c**, N=3. **a-c**, 2-way ANOVA, Bonferroni post-test *p<0.05; ****p<0.0001. **c**, expression of non-FASII genes in WT and mFabT strains (see Supplementary Table 1 for gene assignments). In **a**-**c,** WT, white bars; mFabT, green bars. **a, b**, N=4; **c**, N=3. **a-c**, 2-way ANOVA, Bonferroni post-test *p<0.05;****p<0.0001.

1.Lambert, C. et al. Acyl-AcpB, a FabT corepressor in Streptococcus pyogenes. J Bacteriol 205, e0027423, doi:10.1128/jb.00274-23 (2023).

**Supplementary Fig. 5 | Carbohydrates and amino acid residues produced by human endometrial cells.** Metabolomic analysis of RPMI and HEC-1-A conditioned supernatants, as per legend at right. (see Supplementary Table 5 for complete data). N=3, 2-way ANOVA, Bonferroni post-test; *p<0.05; ***p<0.005; ****p<0.001.

**Supplementary Fig. 6. FabT mutant growth and FA profiles in the absence and presence of exogenous C14:0.** Triplicate cultures of the two isolated *fabT* mutants (FabT^T65M^ and FabT^G99S^) and controls (WT, mFabT, Δ*fabT*) were prepared in BHI medium with 0.025 % BSA. Medium was without (left column), or with 100 µM C14:0 (right column). OD600 is shown for each strain and both conditions after 4 h growth (boxed). For each strain, proportions of C14:0 present in FA profiles of WT and *fabT* mutant strains are in red; proportions, from left to right, of C16:0, C18:0, C18:1Δ9 and C18:1Δ11 are in black.

