## Supplementary Figures 1-6 for "The double-edged role of FASII regulator FabT in *Streptococcus pyogenes* infection"

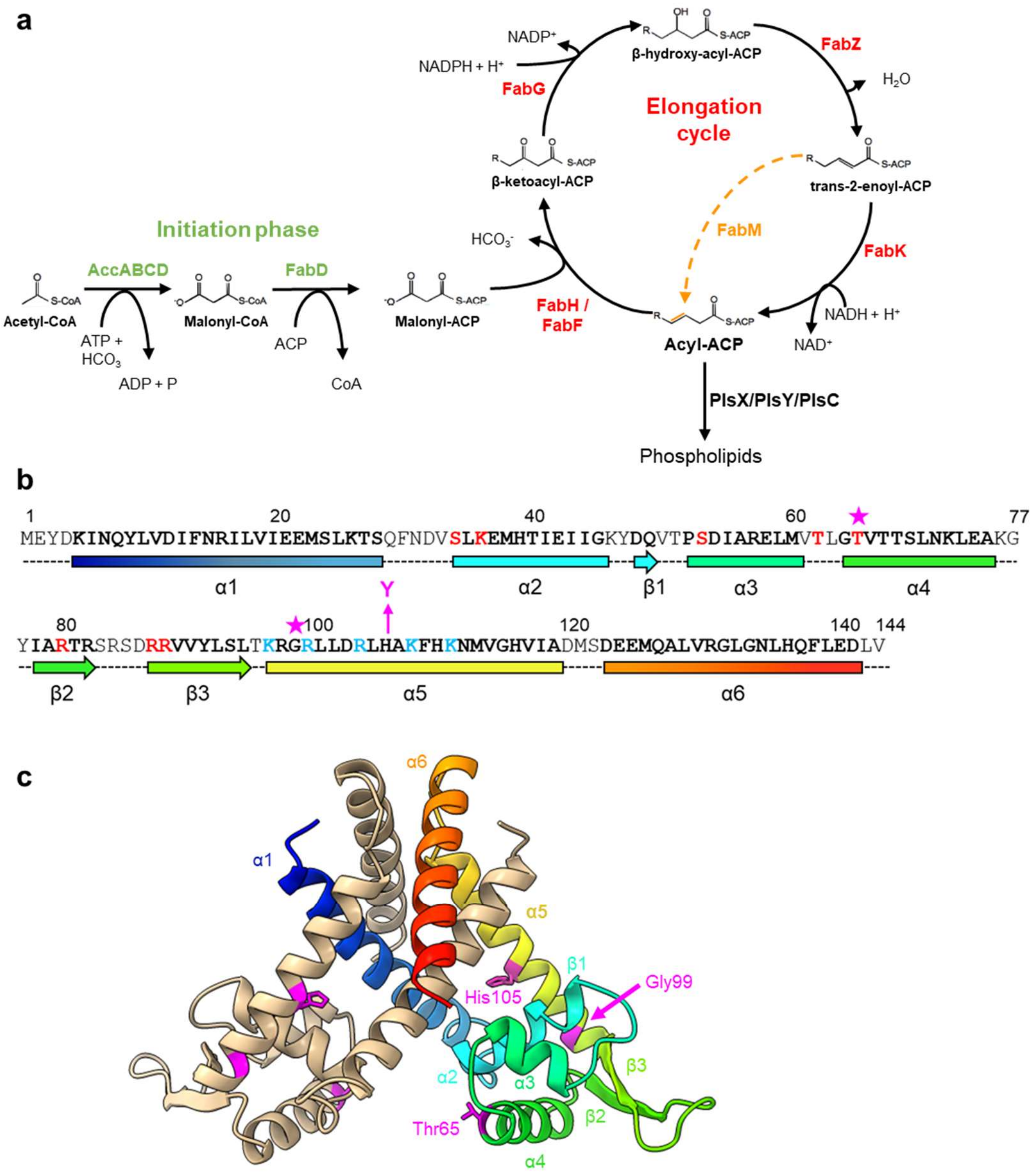


**Supplementary Fig. 1** | **FabT regulator and FASII pathway in GAS. a,** The FASII synthesis pathway comprises a first initiation phase for precursor synthesis, followed by the recursive elongation cycle. The final product, acyl-ACP, supplies FAs for phospholipid synthesis. FabM (orange) leads to unsaturated *cis* FAs; FabK products are saturated. Initiation phase and elongation cycle enzymes are represented in green and red, respectively. From [1]. **b,** FabT sequence; amino acids involved in DNA binding are in red, and those interacting with acyl-Acyl carrier protein are in blue. Arrow indicates the His105Tyr FabT mutation studied in this work. Magenta stars highlight amino acids spontaneously mutated *in vivo* and in a saturated-FA environment (this work). **c,** Overall structure of FabT dimer predicted by Alphafold and adapted with ChimeraX (see references 1-2 in Supplementary Methods); one monomer is represented as multicolored (each color designates a separate domain), and the other is beige. Residues Thr65, Gly99 and His105, in magenta, correspond to mutants isolated in this study.

1. Lambert, C., Poyart, C., Gruss, A. & Fouet, A. FabT, a Bacterial Transcriptional Repressor That Limits Futile Fatty Acid Biosynthesis. *Microbiol Mol Biol Rev*, e0002922, doi:10.1128/mmbr.00029-22 (2022).

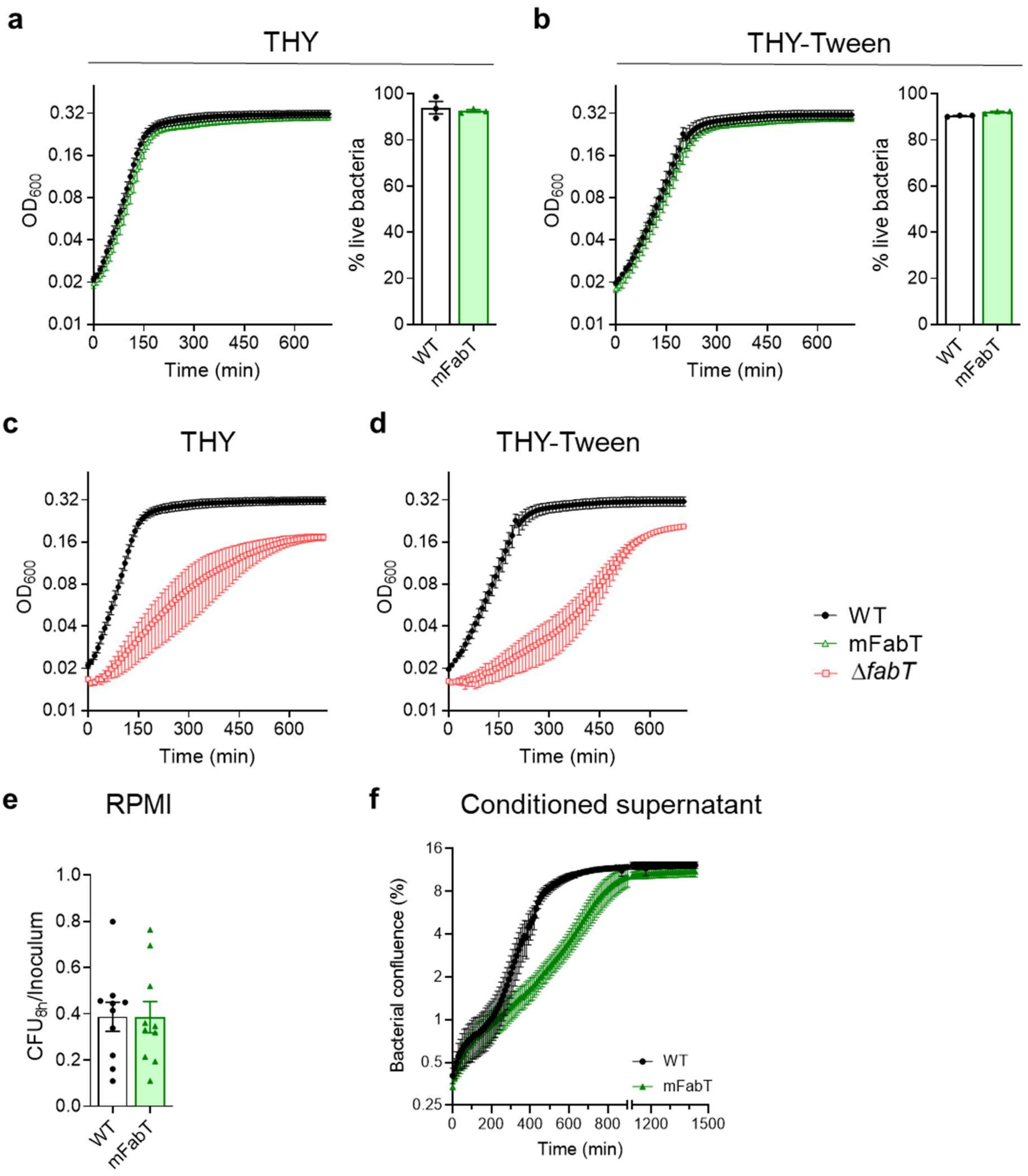


**Supplementary Fig. 2 | Impact of FabT mutations on GAS growth. a-d**, WT growth was compared to mFabT (**a** and **b** left; adapted from reference 1) or to Δ*fabT* (**c** and **d**) on THY and THY-Tween. Viability tests (**a** and **b,** right), using the LIVE/DEAD® BacLightTM Bacterial Viability Kit, were performed on WT and mFabT cultures after growth to $OD_{600}$ = 0.4 - 0.5. Legend at the right of **d** is for growth curves. **e,** Ratios of CFUs of WT or mFabT strains after 8 h over respective initial inocula ($10^3$/ml for each) in RPMI medium; ratio below 1 indicates that bacteria die. **f**, Real-time growth of WT and mFabT in endometrial conditioned supernatant followed using Live-Cell Analysis System (IncuCyte®, Sartorius) and the Basic analyzer. **a-d**, **e**, N=3; **f** N=10; differences in **a, b,** and **e** were not statistically significant using T-test. WT, black lines or white bars; mFabT, green lines or bars. **a, b, e**, Outliers were searched using ROUT method (GraphPad), with Q=1 %. WT, black lines or white bars; mFabT, green lines or bars.


1. Lambert, C. *et al.* Acyl-AcpB, a FabT corepressor in *Streptococcus pyogenes*. *J Bacteriol* **205**, e0027423, doi:10.1128/jb.00274-23 (2023).

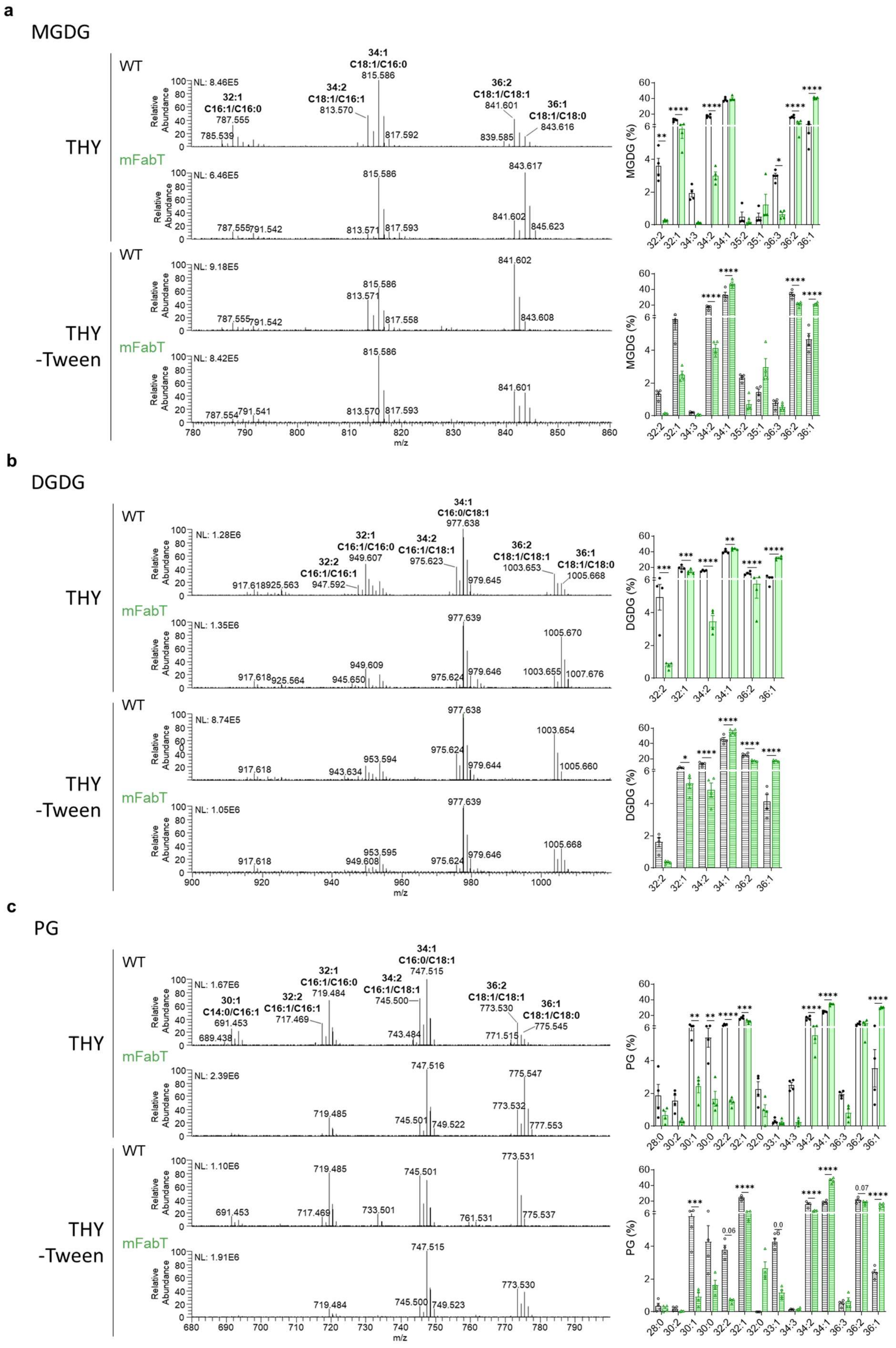


**Supplementary Fig. 3 a-c**

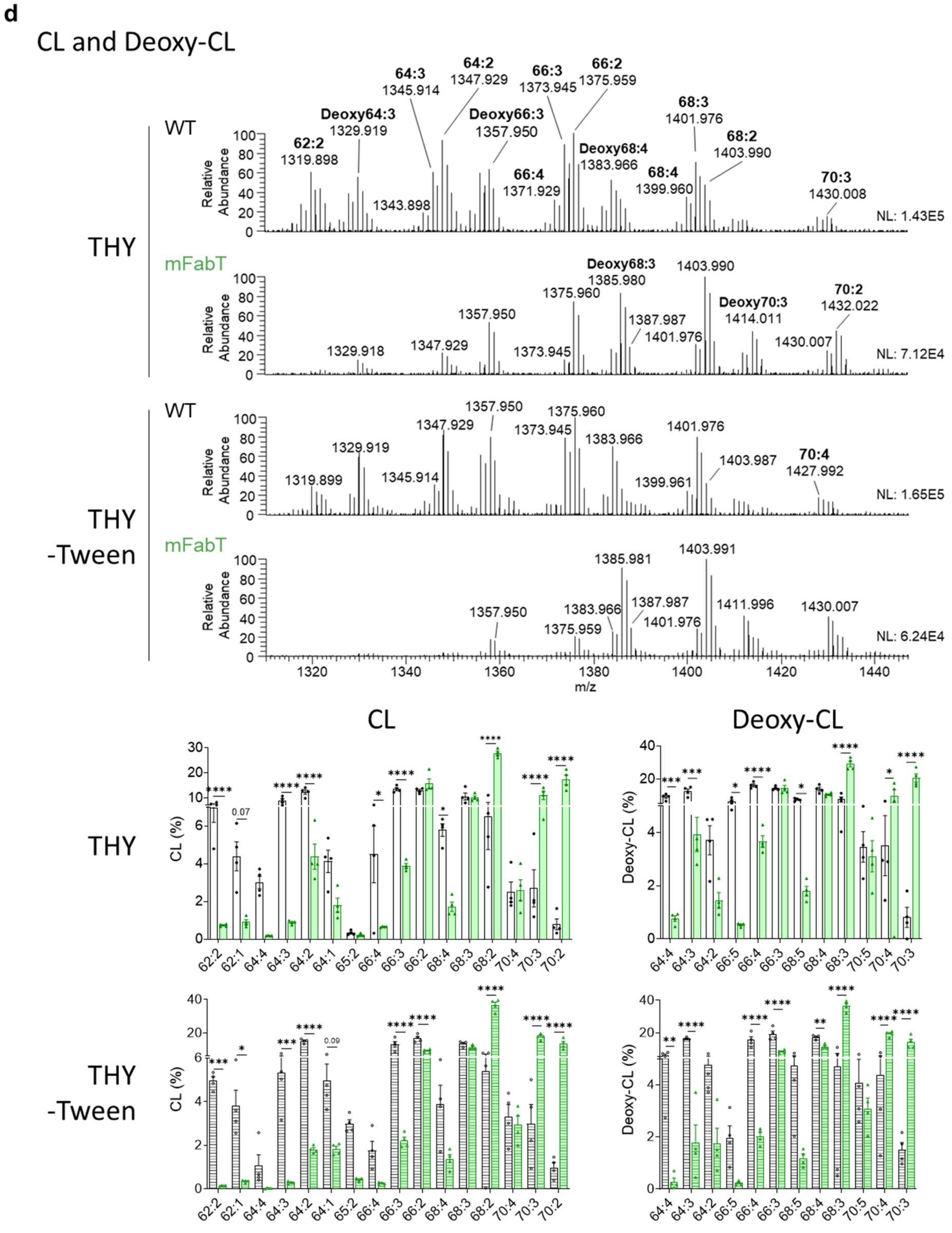


**Supplementary Fig. 3│ Phospholipid membrane composition.** Identification of **a**, monoglucosyldiacylglycerol (MGDG), **b**, diglucosyldiacylglycerol (DGDG), **c**, phosphatidylglycerol (PG), **d**, cardiolipin (CL) and deoxidized cardiolipin (Deoxy-CL). For each class, lipids are presented as the percentage of total lipids, and are quantified in Supplementary Table 2. For **a**, **b**, and **c**, fatty acid assignments are presented directly on mass spectrometry analyses. Cardiolipin assignments were ambiguous and are not shown. Statistical values were determined using 2-way ANOVA, Bonferroni post-test. *p<0.05; **p<0.01; ***p<0.001; ****p<0.0001. Strains were grown in THY (open bars) and THY-Tween (hatched bars). WT, black lines and white bars; mFabT, green lines and bars.

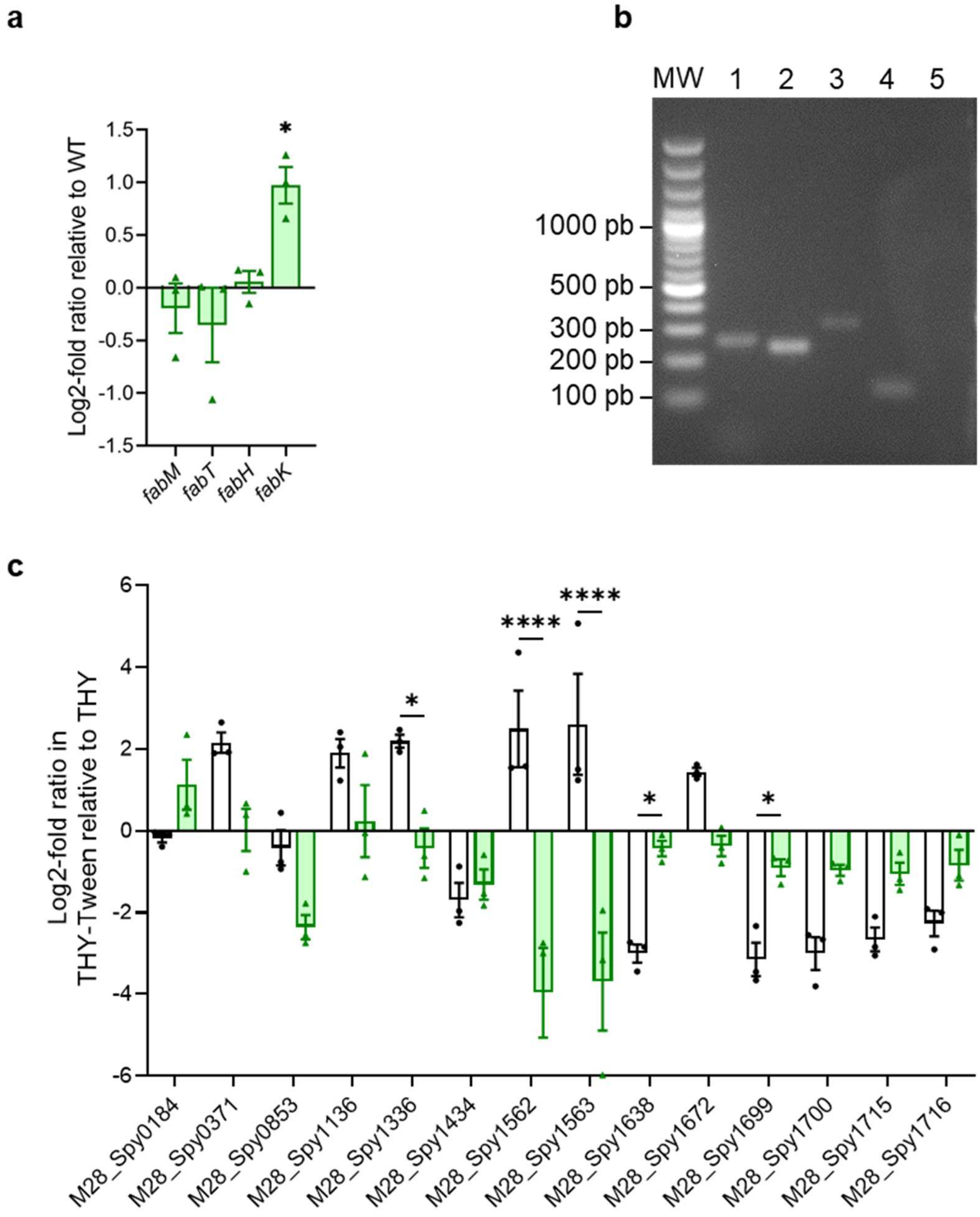


**Supplementary Fig. 4 | Impact of mFabT on FASII and non-FASII gene expression in the absence of FA supplementation**. **a-c**, WT and mFabT strains were grown in THY medium. **a, c**, RNAs were quantified by qRT-PCR. Expression was normalized to that of *gyrA;* relative gene expression is expressed as the log2-fold ratio. **a,** FASII gene expression of mFabT relative to WT is reproduced from Reference 1**. b**, mRNA transcript analysis of FASII locus genes: agarose gel of PCR amplification products on cDNA using primer pairs from neighboring genes. MW, molecular weight reference (Generuler 100 bp, ThermoFisher Scientific). Lane 1, *fabM-fabT* (234 bp); 2, *fabH-acpA* (221 bp); 3, *acpA-fabK* (302 bp); 4, *fabZ-accC* (105 bp); 5, *accD-serS* (288 bp). Results confirm the FASII transcriptional units indicated in Fig. 2a. **c**, expression of non-FASII genes in WT and mFabT strains (see Supplementary Table 1 for gene assignments). The relative gene expression is expressed as the log2-fold ratio in a given strain grown in THY-Tween vs in THY. *, significance of differences in the two media between WT and mFabT. In **a-c,** WT, white bars; mFabT, green bars. **a, b**, N=4; **c**, N=3. **a-c**, 2-way ANOVA, Bonferroni post-test *p<0.05; ****p<0.0001. **c**, expression of non-FASII genes in WT and mFabT strains (see Supplementary Table 1 for gene assignments)**.** In **a-c,** WT, white bars; mFabT, green bars. **a, b**, N=4; **c**, N=3. **a-c**, 2-way ANOVA, Bonferroni post-test *p<0.05; ****p<0.0001.

1. Lambert, C. *et al.* Acyl-AcpB, a FabT corepressor in *Streptococcus pyogenes*. *J Bacteriol* **205**, e0027423, doi:10.1128/jb.00274-23 (2023).

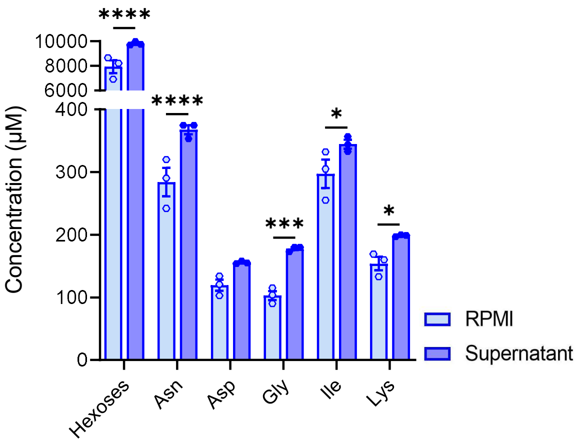


**Supplementary Fig. 5 | Carbohydrates and amino acid residues produced by human endometrial cells.** Metabolomic analysis of RPMI and HEC-1-A conditioned supernatants, as per legend at right. (see Supplementary Table 5 for complete data). N=3, 2-way ANOVA, Bonferroni post-test; *p<0.05; ***p<0.005; ****p<0.001.

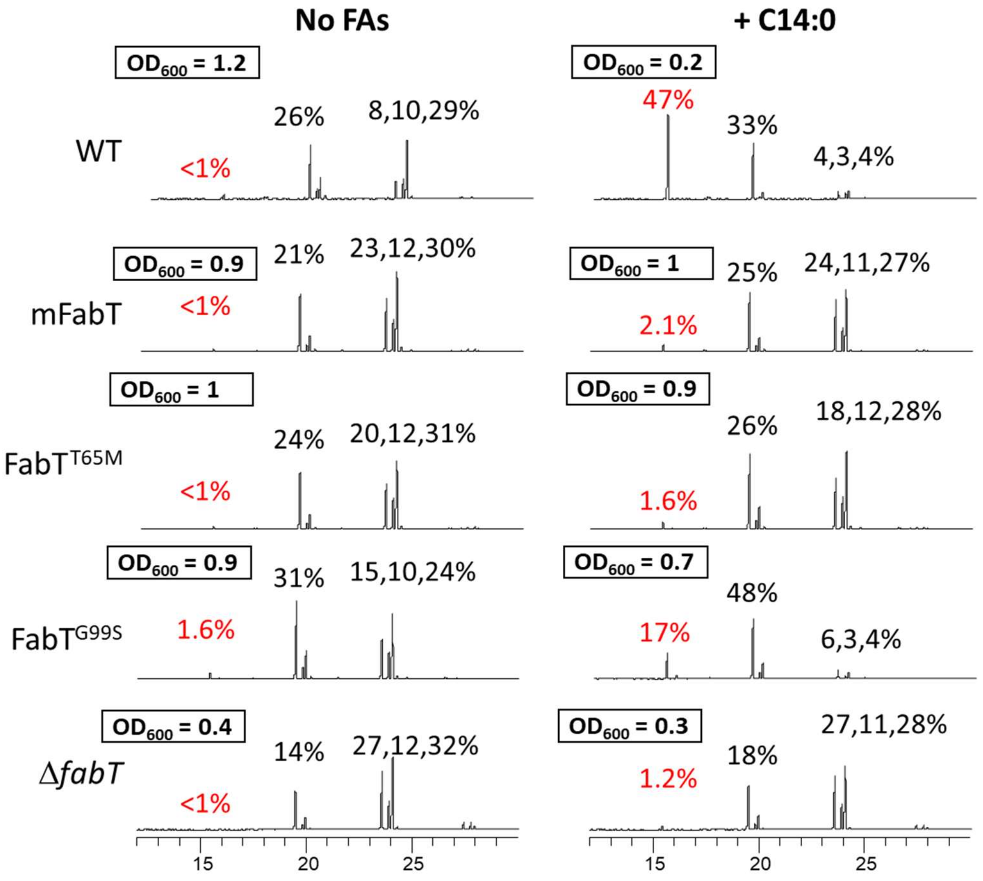


**Supplementary Fig. 6. FabT mutant growth and FA profiles in the absence and presence of exogenous C14:0.** Triplicate cultures of the two isolated *fabT* mutants ($FabT^{T65M}$ and $FabT^{G99S}$) and controls (WT, mFabT, Δ*fabT*) were prepared in BHI medium with 0.025 % BSA. Medium was without (left column), or with 100 µM C14:0 (right column). $OD_{600}$ is shown for each strain and both conditions after 4 h growth (boxed). For each strain, proportions of C14:0 present in FA profiles of WT and *fabT* mutant strains are in red; proportions, from left to right, of C16:0, C18:0, C18:1Δ9 and C18:1Δ11 are in black.
