## Supplementary Methods for "The double-edged role of FASII regulator FabT in *Streptococcus pyogenes* infection"

**FabT modelling**. Overall structure of FabT dimer (supplementary Figure 1) was predicted by Alphafold [1]. We used UCSF ChimeraX [2] for molecular graphics and further analyses. ChimeraX was developed by the Resource for Biocomputing, Visualization, and Informatics at the University of California, San Francisco, with support from NIH R01-GM129325 and the Office of Cyber Infrastructure and Computational Biology, NIAID.

**Live - dead analysis.** Bacterial mortality was determined using the LIVE/DEAD® BacLightTM Bacterial Viability Kit (ThermoFisher Scientific, Ref. L7012) as described for flow cytometry utilization using an ACCURI C6 cytometer (BD Biosciences, Le pont de Claix, France) from the CYBIO Core Facility. Bacteria were grown in HEC-1-A conditioned supernatant for 8 h for testing. Results of three independent experiments were analyzed using the BD Accuri C6 software.

**Real-time bacterial growth**. Test strains were diluted to $10^3$ bacteria per ml in endometrial conditioned supernatant and incubated at 37 °C with 5 % $CO_2$. Bacterial growth was followed by imaging with 20X magnification every 10 min using IncuCyte® Live-Cell Analysis Systems (Sartorius). The bacteria-covered surface was determined using built-in Incucyte software. Experiments were performed in independent triplicates, and visualizations were routinely done on nine positions.

**Growth curves.** GAS stationary precultures were diluted in THY or THY-Tween to an $OD_{600}$ = 0.05, and transferred to 96-well plates, which were incubated at 37 °C in a Thermo

Scientific Multiskan GO (ThermoFischer Scientific). Growth was determined by shaking plates immediately before measuring absorbance at $OD_{600}$ every 10 min.

**GAS growth capacity analysis in RPMI.** GAS bacteria were grown in THY to an $OD_{600}$ = 0.4 to 0.5. Cultures were washed twice in PBS and diluted in RPMI medium without glutamine (Gibco, Ref. 32404-014) to a final concentration of $10^3$ bacteria per ml, and then incubated at 37 °C + 5 % $CO_2$ for 8 h. Serial dilutions were plated on THYA solid medium. Cfus were determined after 24 h growth at 37 °C and normalized to the inoculum for each experiment.

**Screening for FA sensitivity on solid medium.** Overnight WT and mFabT cultures were grown starting from single colonies on solid medium containing 0.5 % bovine serum albumin. Cultures were then adjusted to $OD_{600}$ = 0.1, and 100 µl was spread as a lawn on plates. Four µl (0.1 µm) of each FA (25 mM stocks) were deposited. Plates were incubated 24 h at 37 °C and photographed.

**First-strand cDNA synthesis, quantitative PCR (qRTPCR) and PCR.** First-strand cDNA used for quantitative PCR and PCR experiments was done as follows: 500 nanograms of total RNA was treated with SuperScriptTM II reverse transcriptase and random primers according to manufacturer's instructions (Invitrogen, Life Technologies, France). Quantitative PCR was carried out to determine FASII and non-FASII gene expression with SYBR Green PCR kits (Applied Biosystems, Life Technologies, France**)** as per manufacturer's instructions (Supplementary Table 8). *gyrA* and *rpoB* were used as housekeeping reference genes. Relative quantification of specific gene expression was calculated with the 2−ΔΔCt method using *gyrA* as the reference gene and expressed in log2-fold change. Each assay was performed in triplicate on each sample as described [3]. PCR was carried out for FASII operon mapping using six pairs of primer pairs flanking neighboring FASII genes (Supplementary Table 8) as previously described [4]. PCR amplicons were examined on a 1 % agarose gel.
